# The *Borrelia burgdorferi* adenylyl cyclase, CyaB, is important for virulence factor production and mammalian infection

**DOI:** 10.1101/2021.03.04.433918

**Authors:** Vanessa M. Ante, Lauren C. Farris, Elizabeth P. Saputra, Allie J. Hall, Nathaniel S. O’Bier, Adela S. Oliva Chavez, Richard T. Marconi, Meghan C. Lybecker, Jenny A. Hyde

## Abstract

*Borrelia burgdorferi,* the causative agent of Lyme disease, traverses through vastly distinct environments between the tick vector and the multiple phases of the mammalian infection that requires genetic adaptation for the progression of pathogenesis. Borrelial gene expression is highly responsive to changes in specific environmental signals that initiate the RpoS regulon for mammalian adaptation, but the mechanism(s) for direct detection of environmental cues has yet to be identified. Secondary messenger cyclic adenosine monophosphate (cAMP) produced by adenylate cyclase is responsive to environmental signals, such as carbon source and pH, in many bacterial pathogens to promote virulence by altering gene regulation. *B. burgdorferi* encodes a single non-toxin class IV adenylate cyclase (*bb0723, cyaB*). This study investigates *cyaB* expression along with its influence on borrelial virulence regulation and mammalian infectivity. Expression of *cyaB* was specifically induced with co-incubation of mammalian host cells that was not observed with cultivated tick cells suggesting that *cyaB* expression is influenced by cellular factor(s) unique to mammalian cell lines. The 3’ end of *cyaB* also encodes a small RNA, SR0623, in the same orientation that overlaps with *bb0722*. The differential processing of *cyaB* and SR0623 transcripts may alter the ability to influence function in the form of virulence determinant regulation and infectivity. Two independent *cyaB* deletion B31 strains were generated in 5A4-NP1 and ML23 backgrounds and complemented with the *cyaB* ORF alone that truncates SR0623, *cyaB* with intact SR0623, or *cyaB* with a mutagenized full length SR0623 to evaluate the influence on transcriptional and post-transcriptional regulation of borrelial virulence factors and infectivity. In the absence of *cyaB,* expression and production of *ospC* was significantly reduced, while the protein levels for BosR and DbpA were substantially lower than parental strains. Infectivity studies with both independent *cyaB* mutants demonstrated an attenuated phenotype with reduced colonization of tissues during early disseminated infection. This work suggests that *B. burgdorferi* utilizes *cyaB* and potentially cAMP as a regulatory pathway to modulate borrelial gene expression and protein production to promote borrelial virulence and dissemination in the mammalian host.

## Introduction

*Borrelia burgdorferi*, the causative agent of Lyme disease, is an emerging infectious disease that causes a robust inflammatory multistage disease and accounts for over 80% of all vector-borne illnesses in the United States (Rosenberg 2018; Radolf et al. 2012; Stanek and Strle 2018; Steere et al. 2016). Localized disease presents as flu-like symptoms and is frequently associated with an erythema migrans “bull’s-eye” rash (Steere et al. 2016; Stanek and Strle 2018). If left untreated, the pathogen disseminates to specific tissues with systemic symptoms developing including arthritis, carditis, and encephalomyelitis (Hu 2016; Steere et al. 2016; Stanek and Strle 2018). Patients experience severe morbidity due to ongoing fatigue and malaise as a result of the inflammatory response elicited by *B. burgdorferi*. To date, no human vaccine is available and therapeutics for late stage disease are limited.

*B. burgdorferi* lacks classically-defined virulence factors, such as secretion systems and toxins, and instead relies on dynamic genetic regulation and antigenic variability to invade multiple tissue types and evade the immune system (Radolf et al. 2012; D. Scott Samuels and Samuels 2016). Many studies have noted the responsiveness of *B. burgdorferi* to environmental signals, such as temperature, pH, O_2_, CO_2_, and osmotic stress, as it travels from the tick vector to the mammalian host, but mechanisms of direct environmental detection remain unknown (Carroll, Garon, and Schwan 1999; Carroll, Cordova, and Garon 2000; Popitsch et al. 2017; Konkel and Tilly 2000; Stevenson, Schwan, and Rosa 1995; Tokarz et al. 2004; Seshu, Boylan, Gherardini, et al. 2004; Hyde, Trzeciakowski, and Skare 2007; Bontemps-Gallo, Lawrence, and Gherardini 2016; X. Yang et al. 2000). The BosR-RpoN-Rrp2-RpoS signaling cascade responds to changing environmental cues to allow borrelial adaptation during early mammalian infection and resistance to innate immunity by altering the outer membrane lipoprotein composition (Hyde et al. 2009; Zhiming Ouyang, Deka, and Norgard 2011; Jon S. Blevins et al. 2009; Hyde et al. 2010; Zhiming Ouyang et al. 2009; Z. Ouyang, Blevins, and Norgard 2008; A. H. Smith et al. 2007; M. J. Caimano et al. 2007; 2004; Melissa J. Caimano et al. 2019). Transcription of *rpoS* is regulated by a transcription complex formed of the RNA polymerase, the sigma factor RpoN, the phosphorylated Response regulator protein (Rrp2), and the Borrelia oxidative stress regulator (BosR) (Xiaofeng F. Yang, Alani, and Norgard 2003; Jon S. Blevins et al. 2009; Hyde et al. 2009; Zhiming Ouyang, Deka, and Norgard 2011; Hyde et al. 2010; Zhiming Ouyang et al. 2009; A. H. Smith et al. 2007). The borrelial RpoS regulon includes outer surface lipoproteins DbpA, OspC, and BBK32, and other factors important for tick to mouse transmission and survival in mammalian hosts (He et al. 2007, 32; Hübner et al. 2001; Melissa J. Caimano et al. 2005; 2007; X. F. Yang et al. 2005).

Secondary messengers are a mechanism used by bacterial pathogens, such as *B. burgdorferi*, to modulate gene expression and post-transcriptional regulation in response to environmental signals by altering the function of bound proteins (Yin et al. 2020; Purificação et al. 2020; McDonough and Rodriguez 2011; Savage et al. 2015; Ye et al. 2014; Rogers et al. 2009). In *B. burgdorferi*, the second messenger cyclic di-adenosine monophosphate (c-di-AMP) is essential for in vitro growth and the production of mammalian virulence factors (Ye et al. 2014; Savage et al. 2015). Cyclic di-guanosine monophosphate (c-di-GMP) is a key component of the Hk1-Rrp1 two-component system pathway involved in mammal to tick transmission, midgut survival, motility, and glycerol utilization by *B. burgdorferi* (Zhang et al. 2018; Novak, Sultan, and Motaleb 2014; He et al. 2011; Sultan et al. 2011; Bontemps-Gallo, Lawrence, and Gherardini 2016; Melissa J. Caimano et al. 2015). c-di-GMP is produced by Rrp1 and bound by PlzA to positively regulate glucose metabolism (Rogers et al. 2009; Freedman et al. 2010; Kostick-Dunn et al. 2018; Kostick et al. 2011; Mallory et al. 2016; Sultan et al. 2010; He et al. 2014). Another second messenger cyclic adenosine monophosphate (cAMP) has received less attention in *B. burgdorferi*, but has been found to support virulence in other pathogenic bacteria (McDonough and Rodriguez 2011). cAMP is generated by adenylate cyclases (ACs) to modulate regulation of the bacteria or the host cell depending on which of the 6 classes of AC. cAMP can bind to cAMP receptor proteins (CRP) often resulting in a conformation change that promote efficient binding of specific DNA sites and transcription of numerous genes. *B. burgdorferi* encodes a single class IV AC (*bb0723*), annotated as *cyaB*, which is the smallest of the classes, highly thermostable, and has been identified in only 3 other bacterial species (Khajanchi et al. 2016; Gallagher et al. 2006; N. Smith et al. 2006; Casjens et al. 2000; Dong et al. 2013; Sismeiro et al. 1998). The borrelial genome lacks an annotated CRP or cAMP phosphodiesterase, therefore it is unclear how *B. burgdorferi* generated cAMP might modulate the pathogenesis-specific regulation (Casjens et al. 2000). A previous study confirmed the AC enzymatic activity of recombinant borrelial CyaB (Khajanchi et al. 2016). During infection studies with a transposon *cyaB* mutant strain, it was found that *cyaB* did not play a role in tick to mouse transmission or mammalian infectivity when examined qualitatively by culture outgrowth. A borrelial Tn-seq identified *cyaB* as contributing to resistance to oxidative stress (Ramsey et al. 2017). The overall function of the CyaB enzyme in *B. burgdorferi* signal transduction and virulence factor regulation remains unclear, especially in the absence of any detectable downstream effector molecules that would recognize cAMP.

Borrelial *cyaB* overlaps with an intragenic small RNA (sRNA) SR0623 at the 3’ end of the open reading frame (ORF) and extends into the neighboring *bb0722* gene (Popitsch et al. 2017). sRNAs can be arranged in the genome as antisense, 5’ and 3’ untranslated region (UTR), intergenic, or intragenic (Gottesman and Storz 2011; Papenfort and Vogel 2010; Babitzke et al. 2019). sRNAs have a broad range of function with the ability to regulate translation of target mRNA, degradation of target mRNA, act as a riboswitch, or bind to proteins either altering or sequestering their activity. Recent sRNA transcriptome studies identified over 1,000 putative borrelial sRNA that are regulated in response to temperature shift and nutrient stress (Popitsch et al. 2017; Drecktrah et al. 2018).

In this study, we investigated the role of *cyaB* and SR0623 in borrelial pathogenesis. Our findings indicate that *cyaB* influences mammalian virulence in part through regulation of the BosR-RpoN-Rrp2-RpoS pathway. The regulation of *cyaB* was specific to interactions with host cells, further suggesting this AC is important for the mammalian cycle of pathogenesis and may be responsive to unique host specific signals. CyaB, and possible cAMP signaling, has the potential to be an uncharacterized signaling and regulation pathway important for the progression of Lyme disease.

## Materials and methods

### Growth conditions and media

*E. coli* was grown in Luria-Bertani (LB) broth supplemented with antibiotics at the following concentrations: kanamycin 50µg/ml, spectinomycin 100µg/ml, or gentamicin 15µg/ml. *B. burgdorferi* was grown in Barbour-Stoener-Kelly II (BSKII) medium supplemented with 6% normal rabbit serum (NRS) under microaerophilic conditions at 32°C with 1% CO_2_ unless otherwise stated (Barbour 1984). Modified BSK lacks bovine serum albumin (BSA), pyruvate, and NRS (Ramsey et al. 2017). BSK-lite was made using CMRL 1066 without L-glutamine and without glucose (USBiological) supplemented with 6% NRS and 0.01% L-glutamine (von Lackum and Stevenson 2005). BSK-glycerol is BSK-lite with 0.6% glycerol. BSK media was supplemented with antibiotics at the following concentrations: kanamycin 300µg/ml, streptomycin 100µg/ml, or gentamicin 50µg/ml.

### Plasmid construction and strain generation

Strains, plasmids, and primers generated in this study are listed in Table 1 & S1, respectively. The *cyaB* (*bb0723*) deletion construct was generated by amplifying upstream and downstream regions of approximately 1.5kb and individually TOPO cloned into pCR8 (ThermoFisher) resulting in pJH380 and pJH381, respectively. A KpnI and BamI digest inserted the downstream region from pJH381 into pJH380 to generate pJH383. The P*_flgB_*-*aadA* was PCR amplified and cloned into pCR2.1, designated pJH431, and cloned into pJH383 by SphI and KpnI digest generating the final deletion construct, pJH432. This final construct was transformed into *B. burgdorferi* ML23 and 5A4-NP1, resulting in JH441 and JH522, respectively (Figure 1) (Labandeira-Rey and Skare 2001; Lawrenz et al. 2002). A chromosomal *cyaB* complement construct, pJH446, was generated in the pJH333 backbone that encodes 1.5kb chromosomal regions to allow allelic exchange between *bb0445* and *bb0446,* using P*_flgB_*-*aacC1* as the antibiotic selection (Li et al. 2007; Hyde, Weening, and Skare 2011). JH441 was transformed with pJH446 resulting in strain JH446. The mutant and chromosomal complement strains were transformed with pBBE22*luc* to introduce constitutively expressed bioluminescence (J. S. Blevins et al. 2007; Hyde et al. 2011).

**Figure 1.**
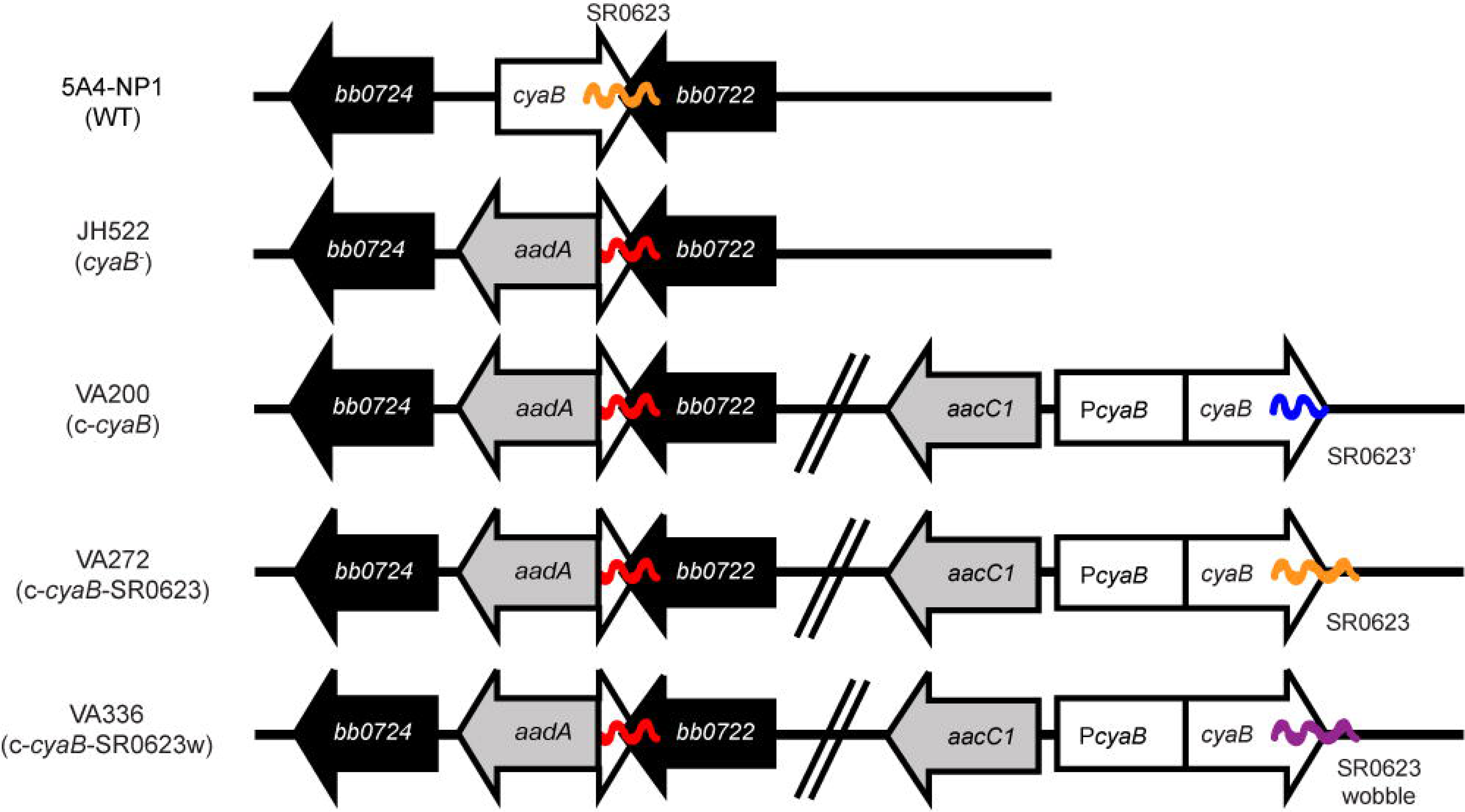
Schematic of the *cyaB* mutant and *trans*-complement strains. The *cyaB* deletion strain JH522 was generated through the insertion of an *aadA* antibiotic cassette to disrupt *cyaB* (*bb0723*) while keeping *bb0722* intact. Chromosomal *cyaB* complementation strains were made through the introduction of an *aacC1* antibiotic cassette. Complement strain VA200 contains the *cyaB* ORF while truncating the sRNA SR0623, strain VA272 contains both the *cyaB* ORF and SR0623, and strain VA336 contains the *cyaB* ORF and a wobble mutation of SR0623. ORFs are indicated by arrows and sRNAs by wavy lines.

**Table 1.**
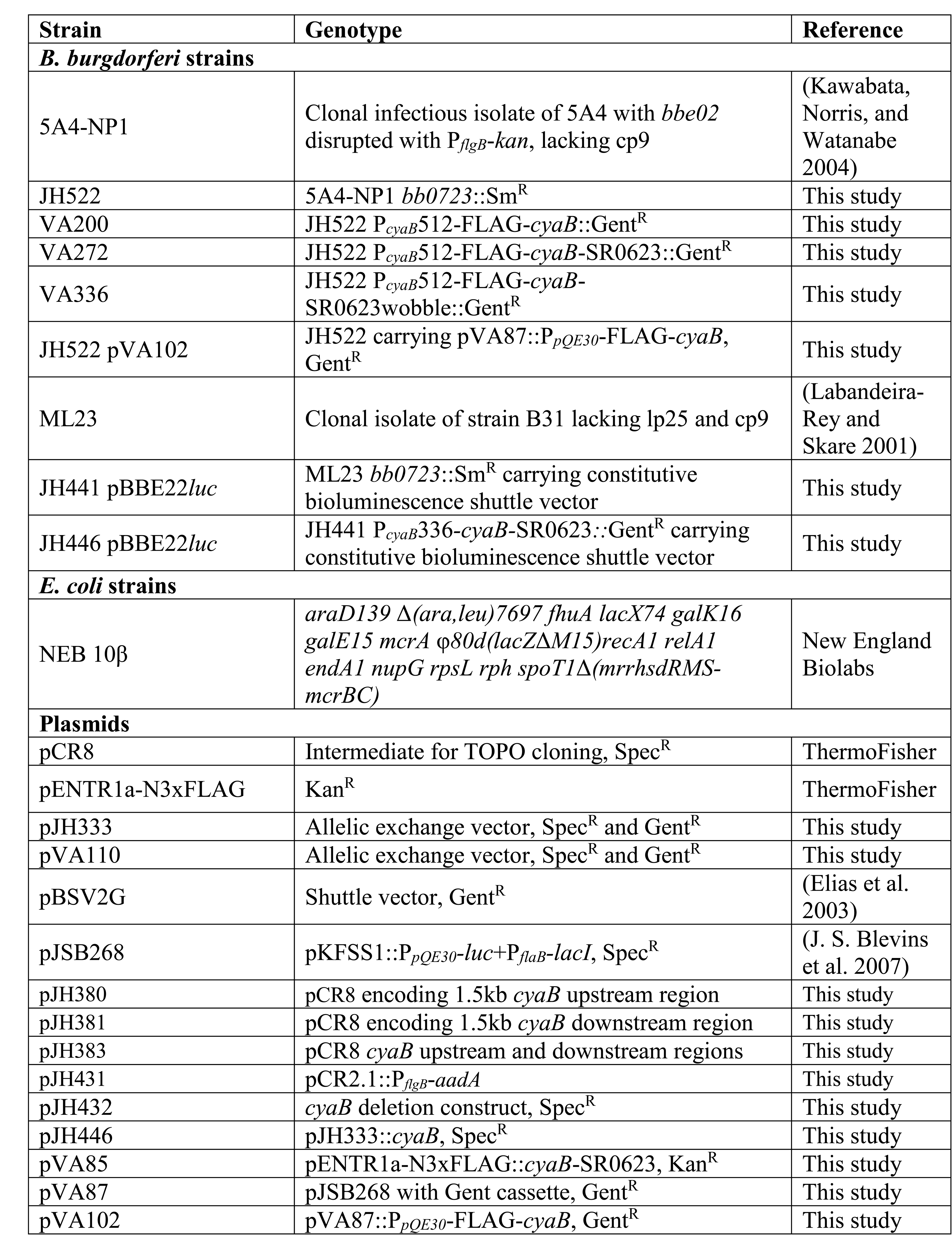
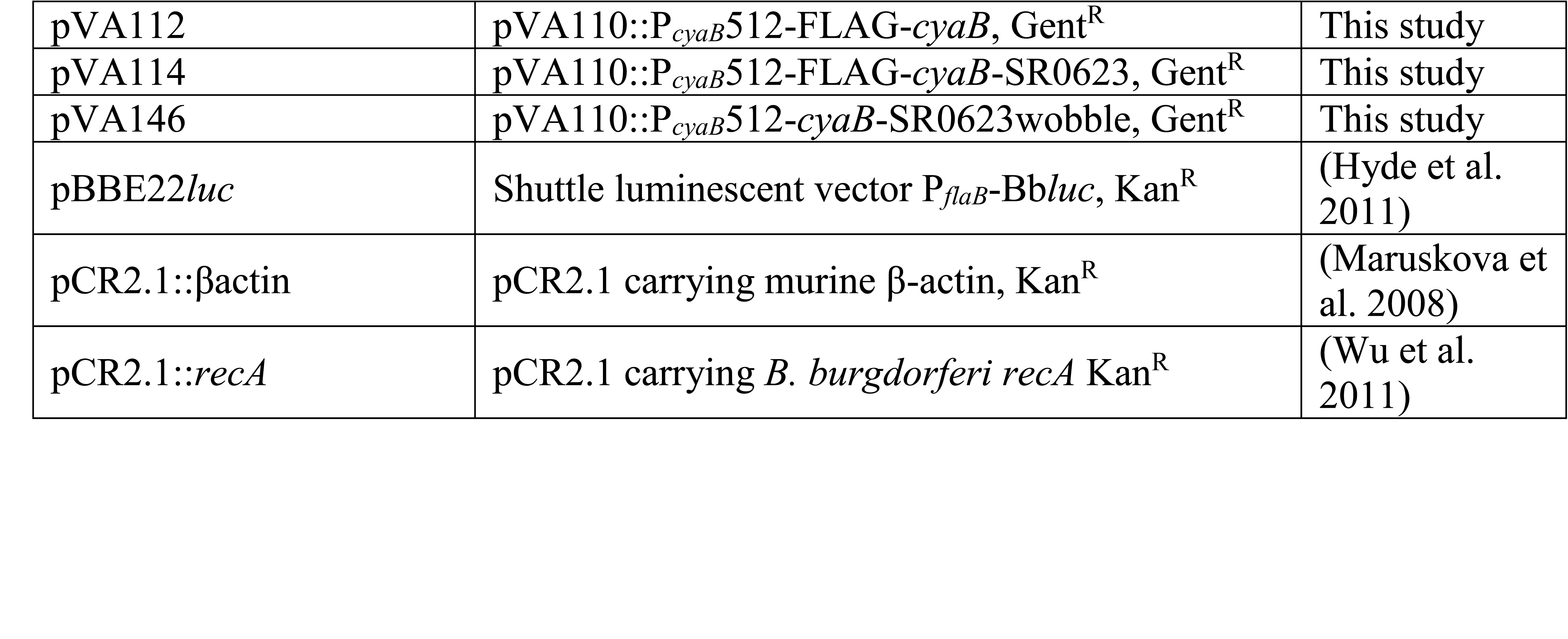
Strains and plasmids used in this study.

A similar construct, pVA110, was generated by amplifying the upstream *bb0445* fragment and P*_flgB_*-*aacC1* using primers *bb0445*-F-BamHI/*bb0445*-R and P*flgB*-F-NotI/gent-R, respectively and underwent overlap PCR with *bb0445*-F-BamHI and P*flgB*-F-NotI, digested with BamHI and NotI, and cloned into pJH333. The *cyaB* complement fragment *cyaB* with SR0623 was PCR amplified using the indicated primers in Table S1, digested with NotI and XhoI, and ligated into pENTR1a-N3xFLAG (ThermoFisher) to create plasmid pVA85. The complement fragment *cyaB* ORF was PCR amplified using primers P*cyaB*-F-SalI/P*cyaB*-R and *cyaB*ORF-F/*cyaB*ORF-R-SalI with pVA85 as the template and underwent overlap PCR with P*cyaB*-F-SalI and *cyaB*ORF-R-SalI. The complement fragment *cyaB* with SR0623 was amplified using primer pair P*cyaB*-F-SalI/P*cyaB*-R and *cyaB*ORF-F/*cyaB*SR0623-R-SalI with pVA85 as the template and underwent overlap PCR with P*cyaB*-F-SalI and *cyaB*SR0623-R-SalI. The modified SR0623 sequence was engineer and manufactured by GenScript to alter the wobble base pair throughout the sRNA (Figure S1). The complement fragments were cloned into pVA110 using the SalI restriction sites resulting in pVA112, pVA114, and pVA146, respectively. Chromosomal complement plasmids were transformed into JH522 generating VA200, VA272, and VA336 (Figure 1 & Table 1).

To make a *trans* inducible FLAG tagged *cyaB* complement with gentamicin resistance for overproduction of CyaB, the NdeI site from P*_flgB_-aacCI* was removed by amplifying the P*_flgB_* and the *aacC1* cassette with pBSV2G as the template. An overlap PCR was performed on the P*_flgB_* and the *aacC1*PCR products, digested, and ligated into pJSB268 digested with AatII and BglII and blunt ended by Klenow (New England Biolabs) to create plasmid pVA87. To generate a N-terminally FLAG tagged *cyaB* construct (pVA102), *cyaB* was amplified from pVA85, digested with NdeI and HindIII, and ligated into pVA87. JH522 was transformed with pVA102.

Electroporation of plasmid DNA into *B. burgdorferi* was done as previously described (D S Samuels, Mach, and Garon 1994; Hyde, Weening, and Skare 2011). Up to 60µg of DNA was transformed, recovered overnight, and then selected for by limiting dilution liquid plating in the appropriate antibiotic and 0.5% phenol red. Transformants were PCR screened for both the allelic exchange and plasmid content (Labandeira-Rey and Skare 2001).

### Oxidative stress assays

Sensitivity to the oxidative stressor H_2_O_2_ was determined as previously performed (Hyde et al. 2009; Ramsey et al. 2017). Briefly, *B. burgdorferi* was grown in BSK-glycerol to mid-log phase, pelleted at 4,800x*g* for 10 min at 4°C, washed with 1X phosphate buffered saline (PBS), and resuspended in modified BSK. 5×10^7^ cells were treated with or without H_2_O_2_, and incubated at 32°C 1% CO_2_ for 4 hours in a 1ml volume. Samples were centrifuged at 6,600xg for 10 min at 4°C, resuspended in BSKII with 6% NRS and 0.6% phenol red, serial diluted in 96-well plates, and incubated for 14 days to assess media color change and survival. Survival was measured from three biological replicates and the data was converted to logarithmic values before calculating the averages.

### *B. burgdorferi* co-cultivation assays

*B. burgdorferi* was co-incubated with *Ixodes scapularis* embryonic cell line ISE6 and human neuroglioma cell line H4 (ATCC HTB-148) to evaluate bacterial transcriptional changes (Schmit, Patton, and Gilmore 2011; J. H. Oliver et al. 1993). ISE6 was maintained in L15C300 supplemented media at 34°C 2% CO_2_ in 25 cm^2^ flasks seeded with 1×10^7^ cells (90% confluency) for 24 hours (J. D. Oliver et al. 2015). H4 cells were seeded at 90% confluency in Dulbecco’s modified Eagle’s medium (DMEM) (Sigma) supplemented with 10% FetalPlex (GeminiBio), herein designated DMEM+, at 37°C 5% CO_2_ for 17 hours. *B. burgdorferi* was grown to mid-log phase under the same conditions as the cell line, cells were pelleted, washed in PBS, and resuspended in cell culture media. *B. burgdorferi* were added to the ISE6 and H4 cells at a multiplicity of infection (MOI) of 10 and 40, respectively (Livengood, Schmit, and Gilmore 2008; J. H. Oliver et al. 1993). Equivalent numbers of borrelial cells were incubated in cell culture media alone as a control. At 3, 6, and 24 hours post infection, the cultivation media was collected for RNA isolation and qRT-PCR analysis.

### Western analysis

*B. burgdorferi* were grown in BSK-lite or BSK-glycerol to mid-log phase and cell lysates were resolved on a 12.5% sodium dodecyl sulfate–polyacrylamide gel electrophoresis (SDS-PAGE), transferred to a polyvinylidene difluoride (PVDF) membrane, and western immunoblotting was conducted as previously described (Labandeira-Rey and Skare 2001; Saputra, Trzeciakowski, and Hyde 2020). Antibody was generated in Sprague-Dawley rats against PlzA as previously described (D. P. Miller et al. 2016; Izac et al. 2019). The following primary antibody concentrations were used: mouse anti-flagellum (1:4000) (Affinity Bioreagent), mouse anti-FLAG (1:4000) (Sigma), rabbit anti-P66 (1:5000) (Cugini et al. 2003), rabbit anti-BosR (1:1000) (Seshu, Boylan, Hyde, et al. 2004), rabbit anti-DbpA (1:10000) (Guo et al. 1998), rat anti-PlzA (1:1000), mouse anti-BadR (1:1000) (C. L. Miller, Karna, and Seshu 2013), mouse anti-OspC (1:20000) (He et al. 2014), mouse-anti-OspA (1:1000) (Capricorn), and mouse anti-Rrp2 (1:1000) (Xiaofeng F. Yang, Alani, and Norgard 2003). Secondary antibodies were coupled to horseradish peroxidase (HRP): donkey-anti-rabbit IgG HRP (Amersham), goat-anti-mouse IgG HRP (ThermoFisher), and rabbit-anti-rat IgG HRP (ThermoFisher). Membranes were imaged with chemiluminescent substrates to detect antigen-antibody complexes. The immunoblot data presented is representative of at least three biological replicates.

### Reverse transcriptase PCR (RT-PCR) and quantitative RT-PCR (qRT-PCR)

*B. burgdorferi* RNA was isolated using hot phenol chloroform extraction as previously described (Meghan Lybecker et al. 2014; M. Lybecker and Henderson 2018). Total RNA was treated with DNaseI (Roche) and 1µg was converted to cDNA using Super Script III reverse transcriptase (+RT) (ThermoFisher) according to the manufacturer’s instructions. A no RT control was included for each RNA sample. PCR reactions using 500ng cDNA as template were amplified with AccuStart II PCR Supermix (Quantabio) and imaged on a 1% agarose gel. qRT-PCR reactions were performed from in vitro cultivated samples with 50ng +RT and –RT cDNA using a ViiA 7 Real-Time PCR system (Applied Biosystems) and Fast SYBR Green Master Mix (Applied Biosystems) according to the manufacturer’s instructions. *B. burgdorferi* co-cultures transcript experiments were performed using PerfeCTa SYBR Green FastMix ROX (Quantabio) and StepOnePlus Real-Time PCR system (Applied Biosystems). *flaB* was used as an internal control and fold change relative to wild type (WT) calculated using the 2^-ΔΔCT^ method from three to four biological and technical replicates (Livak and Schmittgen 2001).

### Northern blots

RNA was collected from *B. burgdorferi* strains grown in BSK-glycerol to mid-log phase at 32°C 1%CO2. RNA isolation and Northern blot analysis was performed as previously described (Popitsch et al. 2017). 7-10 µg of RNA was denatured in 2x RNA load dye (Thermofisher) and heated to 65°C for 15 min, loaded on to a Novex Pre-cast 6% TBE-Urea (8M) polyacrylamide gel (Thermofisher) in 1X TBE and run for 45-60 min. RNA was electroblotted at room temperature (10V for 1 h in 0.5X TBE) to HybondXL membranes (Amersham). The membranes were UV cross-linked (Fisher Scientific UV Crosslinker FB-UVXL-1000) and probed with DNA oligonucleotide (Table S1) in OligoHyb buffer (Thermofisher) per the manufacturer’s protocol. Oligonucleotide probes were end-labeled with γ-^32^P ATP (Perkin-Elmer) and T4 PNK (New England Biolabs) per the manufacturer’s instructions. Unincorporated P^32^ was removed using illustra™ MicroSpin™ G50 columns (GE healthcare). Purified probes were heated at 95°C for 5 min before being added to the prehybridizing bots. Blots were hybridized at 42°C rotating overnight. Membranes were washed 2x 30 min in wash buffer (2x SSC 0.1% SDS). Membranes were placed on Kodiak BioMax maximum sensitivity (MS) autoradiography film and placed in the –80°C for 1-10 days depending on the radiation emission given by each membrane. Film was developed on an AFP imaging developer and scanned using an Epson Expression 10000XL. 5S rRNA was used as the loading control. The Northern blot data presented is representative of three biological replicates.

### Mouse infection studies

Infection studies were conducted using 6-8 week-old C3H/HeN female mice (Charles Rivers) with 5A4-NP1, JH522, VA200, VA272, or VA336. Four to five mice were infected with 10^5^ *B. burgdorferi* by ventral intradermal (ID) injection. Mice were sacrificed and tissues aseptically collected at 7, 14, and 21 days post infection (dpi) for cultivation or qPCR of borrelial load. Outgrowth of viable *B. burgdorferi* was determined by dark-field microscopy and the percent positive tissues was determined. Mice tissues were harvested and DNA was isolated using the DNeasy Blood & Tissue kit (Qiagen) according to the manufacturer’s instructions with the addition 40µl of 10% collagenase (Sigma) and incubated at 55°C overnight. qPCR reactions were performed using a StepOnePlus Real-Time PCR system (Applied Biosystems) and PowerUp SYBR Green Master Mix (Applied Biosystems) according to the manufacturer’s instructions. Standard curves were used to determine the absolute quantification of mouse β-actin and *B. burgdorferi recA*. Technical triplicates were measured for each sample and values are displayed as copies of *B. burgdorferi recA* per 10^6^ mouse β-actin.

To spatially and temporally track luminescent *B. burgdorferi* during infection, an in vivo imaging system (IVIS) was used to image mice (IVIS Spectrum, Perkin Elmer). IVIS infection studies were conducted using 6-8 week-old Balb/c female mice (Charles Rivers) as previously described (Hyde et al. 2011; Hyde and Skare 2018). Briefly, groups of 5 mice were ID infected with 10^5^ *B. burgdorferi* strain ML23 pBBE22*luc*, JH441 pBBE22*luc*, or JH446 pBBE22*luc*. Mice were intraperitoneally (IP) treated with 5mg of D-luciferin and imaged at 1 hour, 1, 4, 7, 10, 14, and 21 dpi. One infected mouse of each group did not receive D-luciferin to serve as a negative control for background luminescence. Images were collected with 1 and 10 min exposures and bioluminescence from the whole body quantitated. Images in the 600-60,000 counts range were used to quantitate bioluminescence. Background bioluminescence was subtracted from the treated samples and averaged. Mice were sacrificed 21 dpi and harvested tissues were used for cultivation as described above.

### Animal ethics statement

Texas A&M University is accredited by the Association for Assessment and Accreditation of Laboratory Animal Care (AAALAC) indicating their commitment to responsible animal care and use. All animal experiments were performed in accordance with the Guide for Care and Use of Laboratory Animals provided by the National Institute of Health (NIH) and the Guidelines of the Approval for Animal Procedures provided by the Institutional Animal Care and Use Committee (IACUC) at Texas A&M University.

### Statistical Analyses

Statistical analysis was performed using GraphPad Prism (GraphPad Software, Inc, La Jolla, CA). The statistical analysis used is listed in the figure legends. Significance was determined by p-values equal to or less than 0.05.

## Results

### Construction of the *cyaB* and SR0623 mutant and complement strains

To investigate *B. burgdorferi cyaB*, we generated a deletion of *bb0723* by replacing the ORF with the P*_flgB_*-*aadA* antibiotic cassette in 5A4-NP1 (WT) resulting in strain JH522 (Figure 1 & Table 1) (Kawabata, Norris, and Watanabe 2004). The 3’ end of *cyaB* overlaps with the 3’ end of its neighboring gene *bb0722* by 26 base pairs, which was not deleted in the *cyaB^-^* strain. In addition, the sRNA SR0623 is encoded within the 3’ end of *cyaB* (95 base pairs) and with *bb0722* (88 base pairs) (Popitsch et al. 2017). SR0623 could either be a result of RNA processing of the *cyaB* mRNA or it could have its own promoter within *cyaB*. Adams et al. globally identified the 5’ end transcriptome and identified putative transcriptional start sites (TSSs) in *B. burgdorferi* (Adams et al. 2017). A putative TSS was not identified within the *cyaB* ORF suggesting SR0623 is synthesized via *cyaB* mRNA processing. To distinguish the functional contribution of CyaB from the sRNA SR0623, we made three chromosomal complements of *cyaB* using its native promoter and P*_flgB_*-*aacC1* antibiotic cassette but included different forms of SR0623 (Figure 1). VA200 encodes *cyaB* and a truncated SR0623, designated c-*cyaB*. A *cyaB* and full length SR0623 is restored in VA272 and named c-*cyaB*-SR0623. A site-directed mutant of SR0623 was generated by altering every third base-pair in the wobble position to disrupt the sRNA primary and secondary structure while maintaining the amino acid sequence of CyaB in complement strain VA336, referred to as c-*cyaB*-SR0623w. For independent verification of the *cyaB* phenotype, deletion and complement strains were also generated in the ML23 background strain. The *cyaB*^-^ strain, JH441, was chromosomally complemented with *cyaB* and SR0623, generating c-*cyaB*-SR0623 strain JH446. JH441 and JH446 were transformed with pBBE22*luc* for in vivo imaging studies (Hyde et al. 2011; Hyde and Skare 2018).

The absence of polar effects on the neighboring genes and confirmation of *cyaB* expression in our strains was verified qualitatively by RT-PCR (Figure 2A). As expected, *cyaB* transcript was detected in WT, c-*cyaB*, c-*cyaB*-SR0623, and c-*cyaB*-SR0623w with a notable absence in the *cyaB^-^* strain. Expression of *bb0722* and *bb0724* was observed in all strains. To evaluate the expression of *cyaB* and SR0623 northern blots were performed with probes designed to hybridize to the 5’ end of SR0623 (Figure 2B). The *cyaB* transcript (484bp) and SR0623 (∼158 bp) are present in the WT strain and absent in the *cyaB^-^* strains, as anticipated. c-*cyaB* produces a *cyaB* transcript and lacks SR0623. c-*cyaB*-SR0623 expresses more *cyaB* and SR0623 than WT strain, which may alter the levels of CyaB protein and perhaps the function of SR0623. c-*cyaB*-SR0623w strain does not have a detectable *cyaB* or SR0623 because the engineered wobble-base mutations prevent the probe from binding. Together, this data suggests that steady-state levels of the *cyaB* mRNA are dependent on its 3’ UTR, which also encodes SR0623.

**Figure 2.**
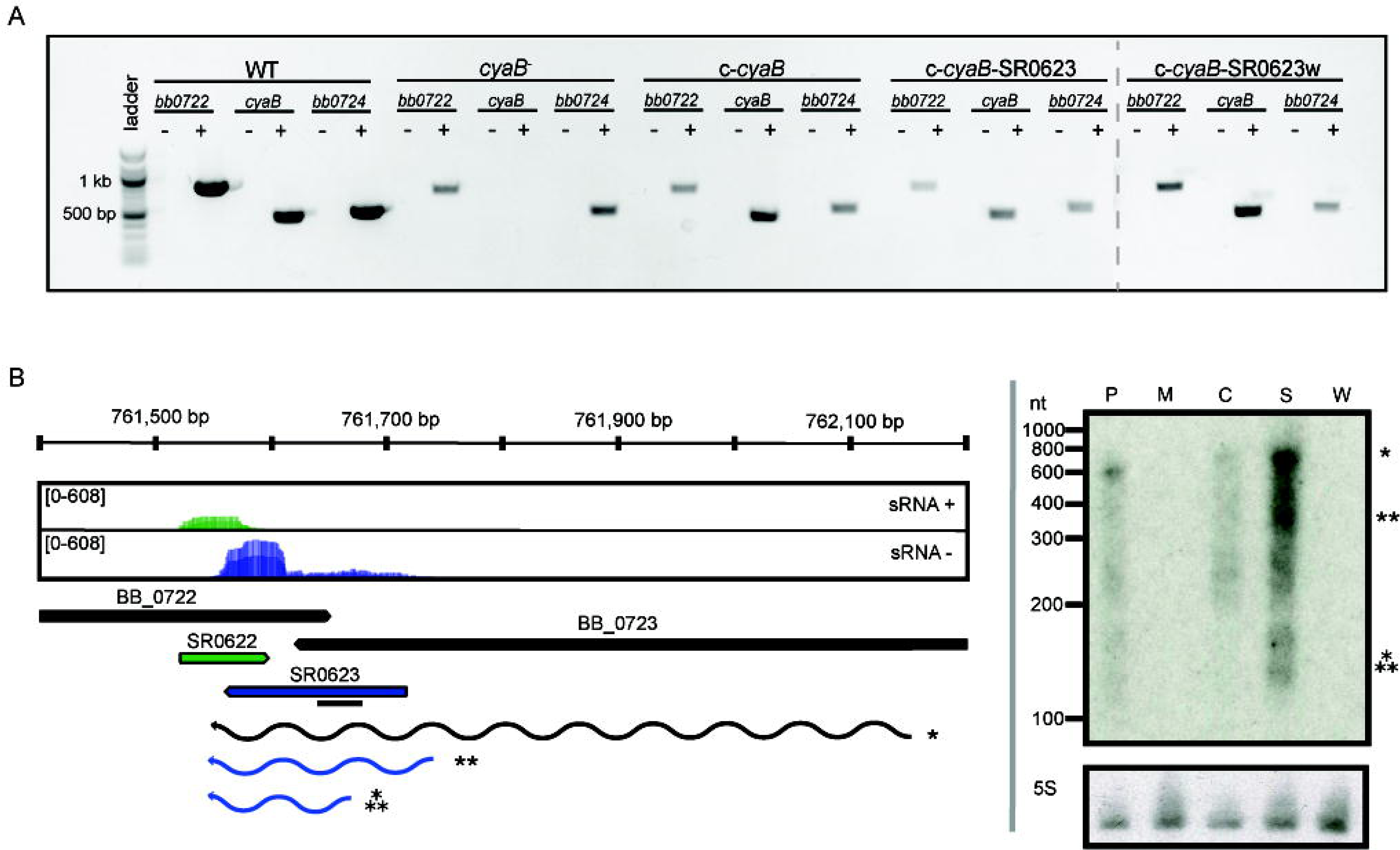
Verification of strains generated in this study. **(A)** Deletion of *cyaB* does not abolish transcription of neighboring genes *bb0722* or *bb0724*. RT-PCR was used to verify the presence or absence of transcripts in the indicated *B. burgdorferi* strains. RNA was isolated and used to generate cDNA as describe in the methods. The +/− symbols indicates if the cDNA reaction with or without reverse transcriptase. PCR reactions included cDNA and primers for *cyaB* (*bb0723*), *bb0722*, or *bb0724*. **(B)** sRNA-sequencing results are displayed in the coverage map (Popitsch et al. 2017). The + strand is shown in green and the – strand in blue. Northern blot analyses of total RNA fractionated on a 6% denaturing polyacrylamide gel, blotted to a nylon membrane and hybridized with radioactive oligonucleotides. The black line represents the location of the oligonucleotide probes. The genomic context is indicated below the coverage plot. The predicted transcripts are denoted and marked with the appropriate band in the Northern blot. Northern blots are representative of three biological replicates. The following abbreviations are used to indicate strains: WT (P), *cyaB*^-^ (M), c-*cyaB* (C), c-*cyaB*-SR0623 (S), c-*cyaB*-SR0623w (W).

### *cyaB* does not contribute to the oxidative stress response

*B. burgdorferi* is able to sense and combat oxidative stress by mechanisms that are not fully understood (Boylan, Posey, and Gherardini 2003; Boylan et al. 2008; Seshu, Boylan, Hyde, et al. 2004; Boylan et al. 2006; Hyde et al. 2009; Ramsey et al. 2017). A Tn-seq screen by Ramsey et al. found *B. burgdorferi* disrupted in *cyaB* had a two-fold decrease in fitness after exposure to H_2_O_2_, therefore we sought out to determine if the AC contributed to the oxidative stress response similar to other pathogens (Ramsey et al. 2017). *B. burgdorferi* strains, WT and *cyaB^-^*, were exposed to increasing concentrations of H_2_O_2,_ then serial diluted to determine the survival percentage. We found that *cyaB^-^* strain survival was comparable to the WT strain 5A4-NP1 (Figure 3). There are several reasons our *cyaB*^-^ phenotype does not correlate with Ramsey et al. *cyaB* Tn-seq phenotype. Ramsey et al. exposed a pool of transposon disruption mutants to an oxidative stressor and measured fitness of the population whereas we are examining a single strain and its survival. Another reason for the difference in findings may be the *B. burgdorferi* growth condition prior to conducting the assay. We grew our cells to mid-log phase in BSK-lite lacking glucose and supplemented with glycerol rather than complete BSKII. It was important for us to limit glucose in the media because in some bacteria, such as *Vibrio cholerae* and *E. coli*, the presence of glucose has been shown to alter AC activity and therefore cAMP production (Liang et al. 2007; Gstrein-Reider and Schweiger 1982). Previous studies used BSK-glycerol to grow *B. burgdorferi* when investigation of secondary metabolite c-di-GMP was evaluated (He et al. 2011). We found that absence or addition of glycerol to the BSK-lite did not influence *B. burgdorferi* growth or alter mammalian virulence factor production (Figure S2).

**Figure 3.**
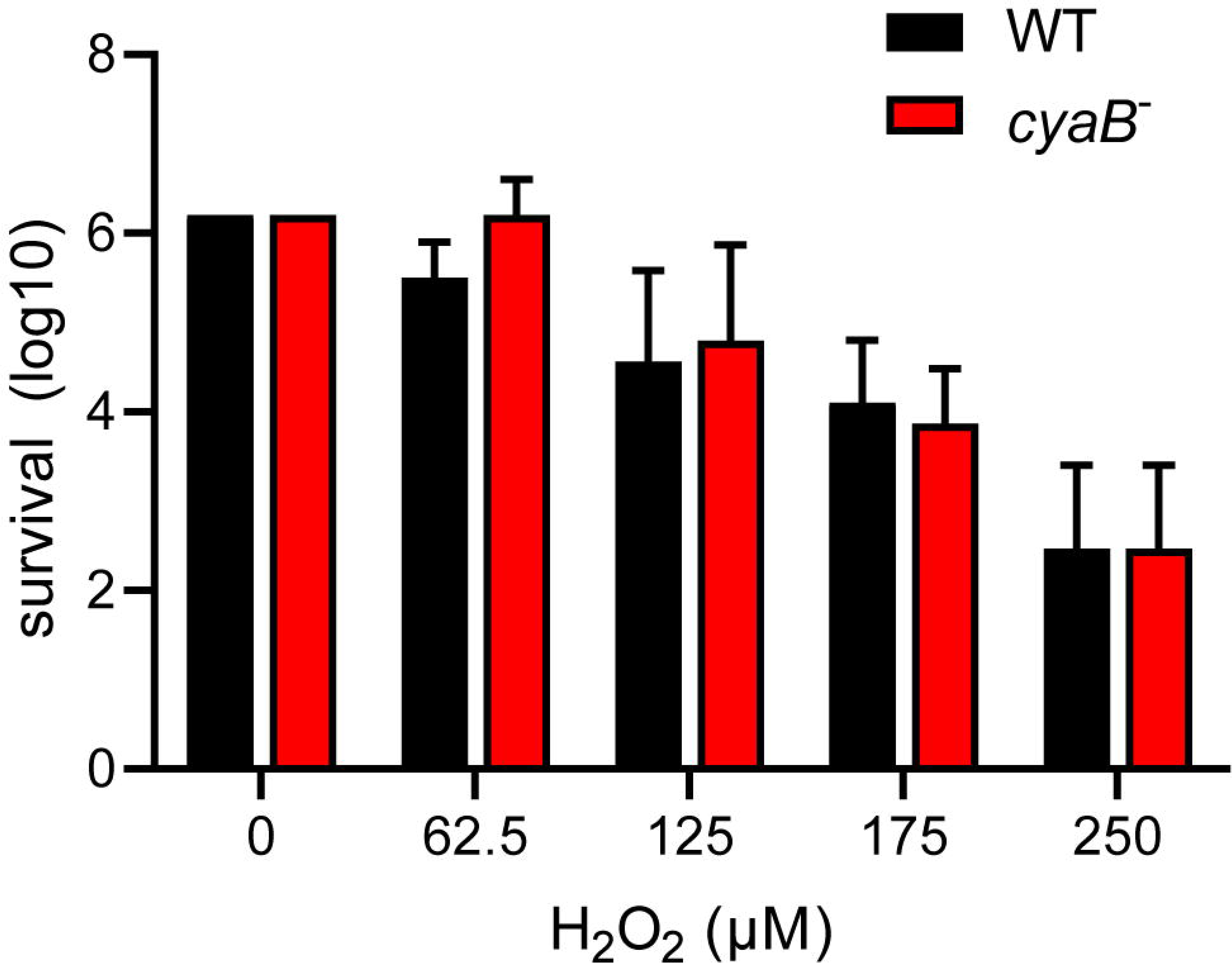
*cyaB* does not contribute to H_2_O_2_ resistance. WT and *cyaB*^-^ strains were grown in BSK-glycerol to mid-log phase at 32°C 1% CO_2_, exposed to H_2_O_2_ in modified BSK media for 4 hours at 32°C 1% CO_2_, and serial diluted for outgrowth to determine survival. Shown is the average and standard error of three independent biological replicates.

### *cyaB* influences borrelial virulence determinants

Loss of an AC results in attenuation of virulence factors in many different bacteria, such as *Pseudomonas aeruginosa* and *Salmonella typhimurium,* therefore we were interested in examining the influence of *cyaB* on important *B. burgdorferi* virulence determinants (R. S. Smith, Wolfgang, and Lory 2004; Curtiss and Kelly 1987). Transcript levels were measured for multiple *B. burgdorferi* targets that included genes important for regulation in the tick vector (*hk1*, *rrp1*, *plzA*, and *ospA*) and mammalian virulence genes (*badR*, *bosR*, *rpoS*, *ospC*, *dbpA*, and *bbk32*) (Figure 4). No changes were observed for genes shown to be operative in the arthropod-borne phase of infection, which may be due to secondary messenger c-di-GMP being involved in the vector (Figure 4A) (Rogers et al. 2009; He et al. 2014). Surprisingly, only *ospC* transcription was found to be statistically significantly reduced 14-fold in the *cyaB^-^* strain compared to WT (Figure 4B). The expression of *ospC* was fully and partially restored to WT level in the c-*cyaB*-SR0623w and c-*cyaB* strains, respectively. The c-*cyaB*-SR0623 had an *ospC* transcript level similar to the *cyaB*^-^ strain. This transcriptional data indicates that *cyaB* may be able to influence *B. burgdorferi* mammalian virulence determinant expression.

**Figure 4.**
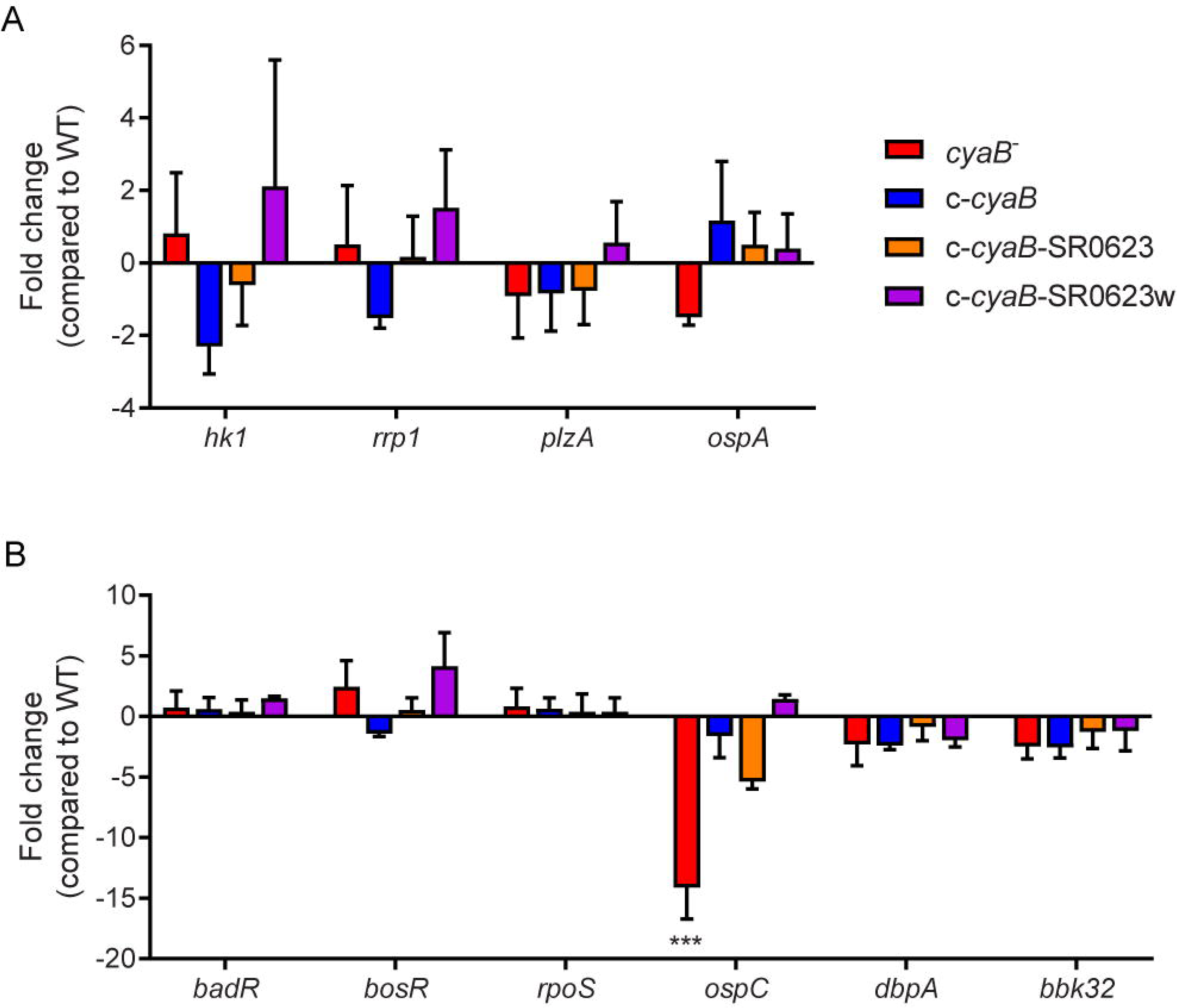
Deletion of *cyaB* reduces *ospC* expression. *B. burgdorferi* was grown in BSK-glycerol to mid-logarithmic phase at 32°C 1% CO_2_ for qRT-PCR analysis to investigate the relative transcript levels of virulence determinants for the **(A)** tick and **(B)** mammalian cycle. *flaB* was used as an endogenous control and fold changes are compared to WT. Shown are the averages and standard error of three biological replicates. Statistical analysis was performed using one-way ANOVA with Dunnett correction relative to WT, *** p<0.001.

cAMP plays an important role in post-transcriptional regulation in many bacteria, for example, in *S. enterica* cAMP-CRP post-transcriptionally regulates transcriptional regulator HilD resulting in reduced virulence factor production (Mouali et al. 2018). Our next step was to evaluate the impact of *cyaB* on the protein production of *B. burgdorferi* virulence determinants. We used immunoblotting to examine the protein levels of several borrelial components associated with the tick and mammalian pathways (Figure 5). Virulence determinants PlzA and OspA, important for survival in the tick, and mammalian virulence determinants Rrp2 and BadR, were not altered in the *cyaB*^-^ strain compared to WT. *bosR* undergoes transcriptional and post-transcriptional regulation in response to pH or metals and CO_2_, respectively, by unknown mechanisms, therefore we considered that BosR may also be post-transcriptionally regulated by cAMP (Hyde, Trzeciakowski, and Skare 2007; Saputra, Trzeciakowski, and Hyde 2020). We found that the *cyaB*^-^ strain produced less BosR relative to WT demonstrating another condition where BosR is post-transcriptionally regulated given there was no difference observed in *bosR* expression (Figure 4 & 5). Strains c-*cyaB* and c-*cyaB*-SR0623w restored BosR protein production back to approximate WT levels. However, the c-*cyaB*-SR0623 complement had BosR protein levels comparable to the *cyaB^-^* strain. The lack of complementation of c-*cyaB*-SR0623 may be in part due to different expression levels of *cyaB* and SR0623 observed by Northern analysis (Figure 2B). DbpA and OspC protein production followed the same pattern as BosR. The different phenotypes of complement *cyaB* strains suggests a possible regulation of *cyaB* by SR0623. Collectively, these results would indicate that CyaB or cAMP, directly or indirectly, post-transcriptionally activates both BosR and DbpA. This data confirms that the changes observed in the *ospC* transcript influence the OspC protein level. It remains unclear if OspC and DbpA are being regulated by *cyaB* independently or through BosR regulation of *rpoS*.

**Figure 5.**
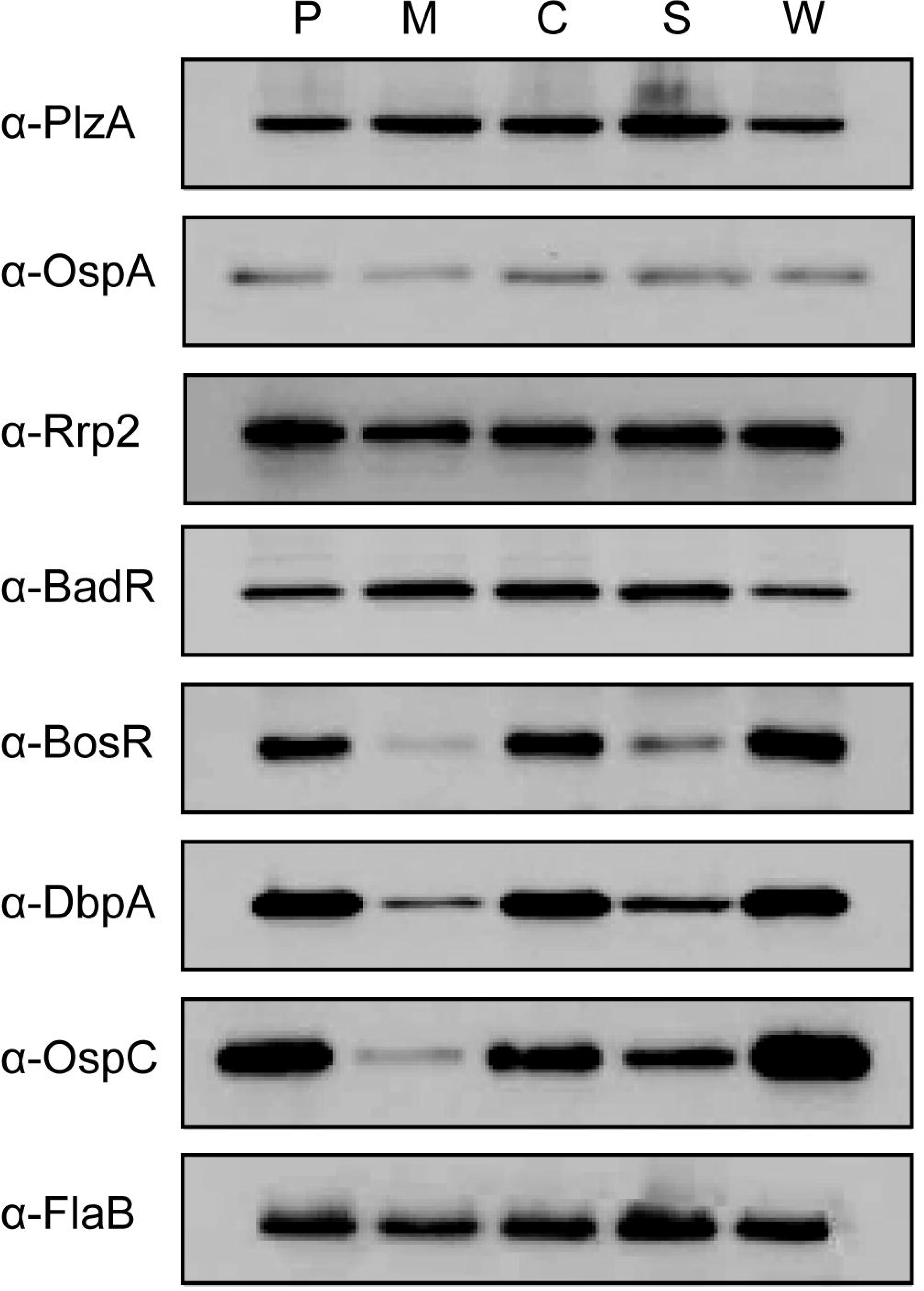
Deletion of *cyaB* reduces protein production of BosR, OspC, and DbpA. *B. burgdorferi* was grown in BSK-glycerol to mid-logarithmic phase at 32°C 1% CO_2_. Protein was harvested and resolved on a SDS-PAGE with approximately 4×10^7^ *B. burgdorferi* in each lane. Immunoblots were prepared using the depicted anti-serum. FlaB was used as a loading control. Displayed is representative of three independent replicates. The following abbreviations are used to indicate strains: WT (P), *cyaB*^-^ (M), c-*cyaB* (C), c-*cyaB*-SR0623 (S), c-*cyaB*-SR0623w (W).

### *cyaB* expression in induced by host cell interaction

*B. burgdorferi* gene expression is highly responsive to changes in various environmental cues and may also impact *cyaB* expression. Under BSK cultivation with shifts in temperature, pH, and CO_2_ we observed no significant changes in *cyaB* transcript (data not shown). To investigate if expression of *cyaB* is influenced by tick or mammalian cellular factors, we co-cultured *B. burgdorferi* strain 5A4-NP1 for 24 hours with the tick neuroglial cell line ISE6 or the mammalian neuroglial cell line H4 and harvested bacteria in the cell culture media (Figure 6) (Livengood, Schmit, and Gilmore 2008; Oliver et al. 1993). *cyaB* expression was not significantly changed at the time points tested in the tick neuroglial cell line ISE6 relative to *B. burgdorferi* incubated in cell culture media alone. Mammalian neuroglial cell line H4 significantly induced the borrelial *cyaB* expression after 24-hour co-incubation. These results indicate that *cyaB* expression is influenced by cellular factor(s) unique to mammalian cell lines and not by in vitro grown tick cells or during in vitro cultivation in BSKII media.

**Figure 6.**
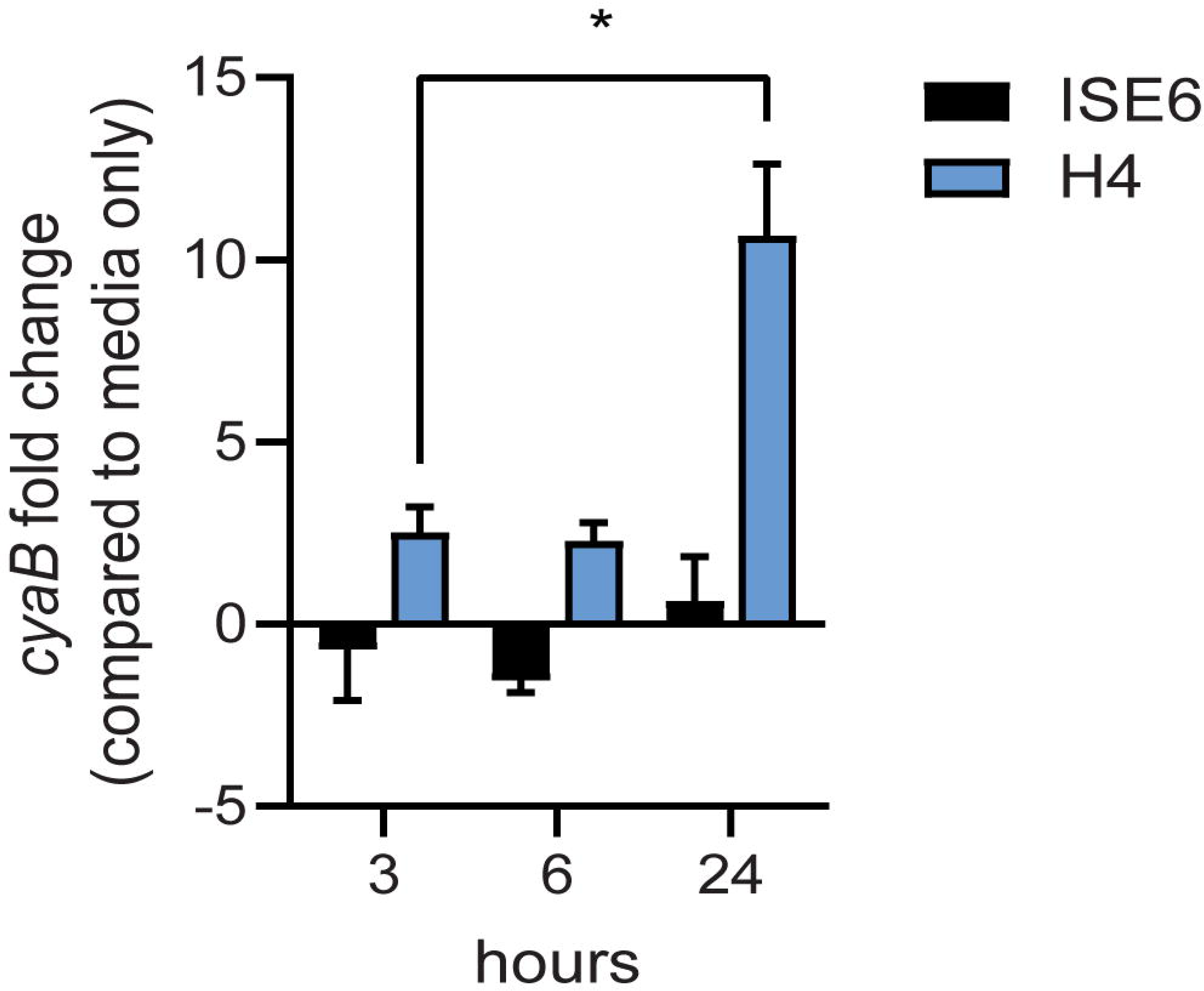
*cyaB* expression is induced with mammalian H4 cell co-culture. *B. burgdorferi* was co-cultured with ISE6 tick cells or H4 mammalian cells. qRT-PCR was performed on samples collected at 3, 6, and 24 hours co-incubation. *flaB* was used as an endogenous control. Shown is the average and standard error of four biological replicates. Statistical analysis was done using Two-way ANOVA with Tukey correction, *p-value<0.05.

### The absence *cyaB* results in attenuated infection

Given that deletion of *cyaB* resulted in altered mammalian virulence determinants, we hypothesized that infecting mice with the *cyaB*^-^ strain would alter infectivity. C3H/HeN mice were intradermally infected with the WT, *cyaB*^-^ strain, or complement strains. Tissues were collected at 7, 14, and 21 days post-infection (dpi) for both qualitative outgrowth and quantitative molecular analysis of infection. Cultivation data shows that all harvested tissues from mice infected with the WT strain were positive for *B. burgdorferi* by day 14, whereas, half the tissues for the *cyaB^-^* strain did not have bacterial outgrowth (Table 2). Unfortunately, none of the three complement strains were able to restore infectivity to WT levels. This was surprising given that the qRT-PCR, Northern, and Western data show that the c-*cyaB* and c-*cyaB*-SR0623w strains restore the *cyaB* in vitro deletion phenotypes. *B. burgdorferi* burden of individual infected tissues was analyzed by qPCR to determine the bacterial burden. The *cyaB^-^* strain was significantly lower at 7 dpi in lymph nodes, bladders, and skin flanks adjacent to the inoculum site. The ears, a distal skin colonization, and joints had less *B. burgdorferi* at 14 and 21dpi, respectively, when infected with the *cyaB^-^* strain (Figure 7). The *cyaB* complements had borrelial loads similar to the *cyaB^-^* strain. Taken together this data would indicate that *cyaB* plays a role in mammalian infectivity.

**Figure 7.**
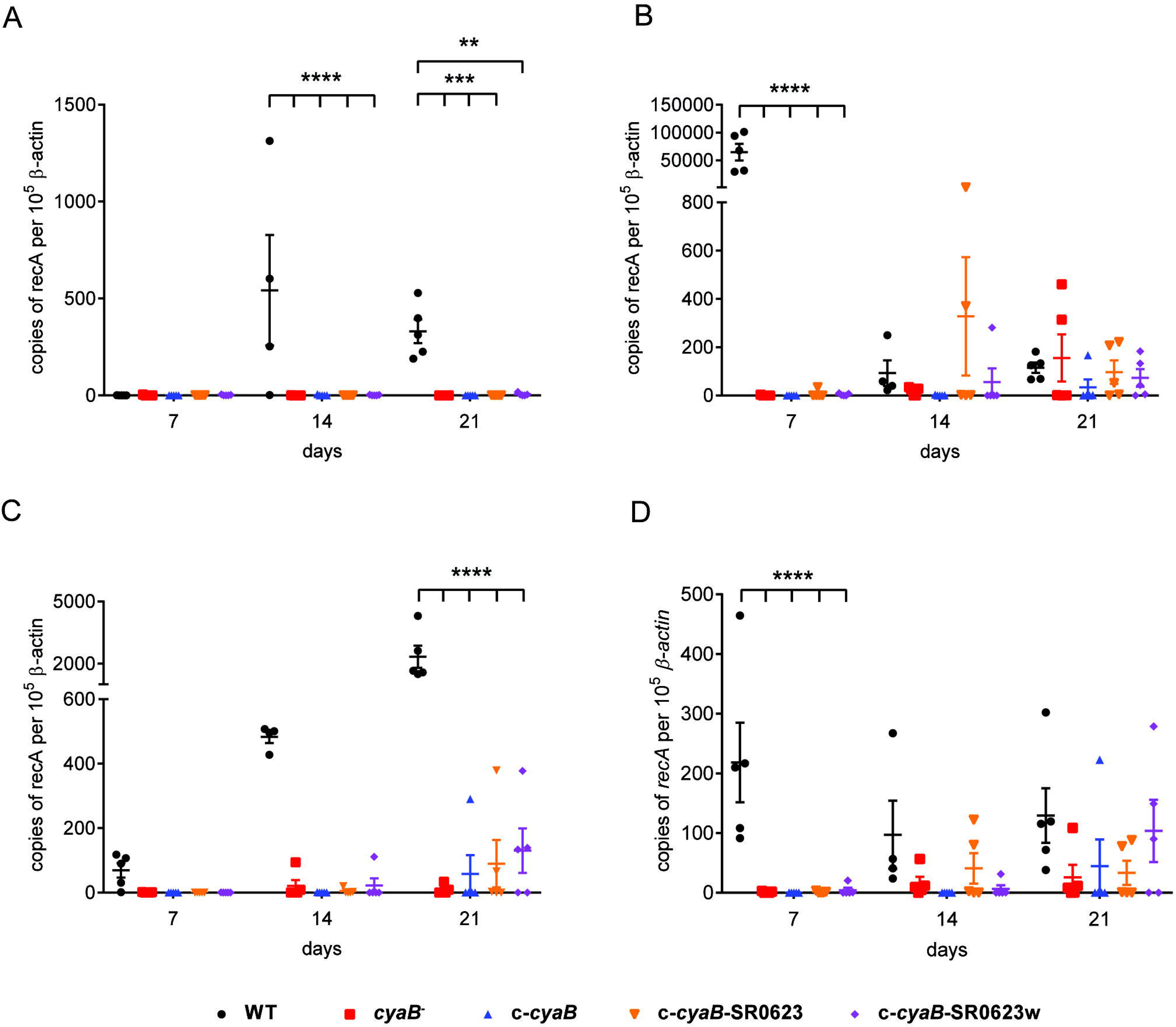
Borrelial burden and tissue dissemination is reduced in mouse tissues infected with the *cyaB* mutant. C3H/HeN mice were infected with 10^5^ *B. burgdorferi* and tissues were harvested at 7, 14, and 21 dpi. qPCR was performed on **(A)** Ears, **(B)** Skin Flanks, **(C)** Joints, and **(D)** Bladders to determine the number of borrelial genomes (*recA*) per 10^6^ copies of mouse β-actin. Individual data points with at least n of 4 with lines representing average and standard error. Statistical analysis was done using Two-way ANOVA with Dunnett correction relative to WT, **p-value<0.01, ***p-value<0.001, ****p-value<0.0001.

**Table 2.** Tissue infectivity of *B. burgdorferi* infected mice.

| **Day 7** | number of positive cultures/total |  |  |  |  |
| --- | --- | --- | --- | --- | --- |
| strain | lymph node | skin flank | ear | all sites | % positive all sites |
| WT | 5/5 | 5/5 | 1/5 | 11/15 | 73 |
| *cyaB*<sup>-</sup> | 0/5 | 1/5 | 0/5 | 1/15 | 6 |
| *c-cyaB* | 0/5 | 0/5 | 0/5 | 0/15 | 0 |
| *c-cyaB*-SR0623 | 1/5 | 1/5 | 0/5 | 2/15 | 13 |
| *c-cyaB*-SR0623w | 2/5 | 4/5 | 0/5 | 6/15 | 40 |

| **Day 14** | number of positive cultures/total |  |  |  |  |
| --- | --- | --- | --- | --- | --- |
| strain | lymph node | skin flank | ear | all sites | % positive all sites |
| WT | 4/4 | 4/4 | 4/4 | 12/12 | 100 |
| *cyaB*<sup>-</sup> | 3/5 | 2/5 | 0/5 | 5/15 | 33 |
| *c-cyaB* | 0/5 | 0/5 | 0/5 | 0/20 | 0 |
| *c-cyaB*-SR0623 | 2/5 | 2/5 | 0/5 | 4/15 | 26 |
| *c-cyaB*-SR0623w | 1/5 | 1/5 | 1/5 | 3/15 | 20 |

| **Day 21** | number of positive cultures/total |  |  |  |  |
| --- | --- | --- | --- | --- | --- |
| strain | lymph node | skin flank | ear | all sites | % positive all sites |
| WT | 5/5 | 5/5 | 5/5 | 15/15 | 100 |
| *cyaB*<sup>-</sup> | 2/5 | 2/5 | 0/5 | 4/15 | 26 |
| *c-cyaB* | 1/5 | 1/5 | 0/5 | 2/15 | 13 |
| *c-cyaB*-SR0623 | 3/5 | 3/5 | 0/5 | 6/15 | 40 |
| *c-cyaB*-SR0623w | 3/5 | 3/5 | 1/5 | 7/15 | 46 |

To independently evaluate the influence of *cyaB* on infectivity, we generated a *cyaB* deletion in the ML23 background that is used to track bioluminescent imaging during murine infection as a way to evaluate temporal and spatial dissemination (Hyde et al. 2011, 32; Hyde and Skare 2018). The parent (ML23), *cyaB*^-^ (JH441), and c-*cyaB-*SR0623 complement (JH446) strains had pBBE22*luc* introduced into them were tested for disseminated infectivity using light emission as a reporter for live *B. burgdorferi* (Hyde et al. 2011). Western analysis of the aforementioned showed less protein production of BosR, DbpA, and OspC in the *cyaB*^-^ strain as was observed in the 5A4-NP1 background mutant (Figure S3). Balb/c mice were then infected with the *B. burgdorferi* strains and in vivo imaged at 0, 1, 4, 7, 10, 14, and 21 dpi. The bioluminescent tracking shows the bacteria being localized to the site of injection early at day 0 and then progressing to distal tissues throughout the mouse by day 21 (Figure 8A). At 7 dpi, the peak of infection, the parent strain produces three times more light than the *cyaB*^-^ strain; however, the c-*cyaB-*SR0623 complement strain is not able to restore the light emission observed for the parent strain (Figure 8B). The parent strain disseminates to distal tissues through day 21. This dissemination is not observed in the *cyaB*^-^ or c-*cyaB-*SR0623 strains, which instead stay localized near the site of injection. After imaging on day 21, mice tissues, lymph nodes, skin flanks, ears, and joints, were harvested and used for cultivation. We found a slight reduction in the number of infected tissues in *cyaB*^-^ strain and c-*cyaB-*SR0623 *B. burgdorferi* (Figure 8C). It is interesting to note that the ears of the *cyaB*^-^ strain and c-*cyaB*-SR0623 strain infected mice were negative for *B. burgdorferi*, suggesting the genetic modifications may alter tissue dissemination. This independently validates the infectivity data in the 5A4-NP1 background and, taken together, despite the issues with incomplete complementation, supports the finding that *cyaB* is important for murine infection.

**Figure 8.**
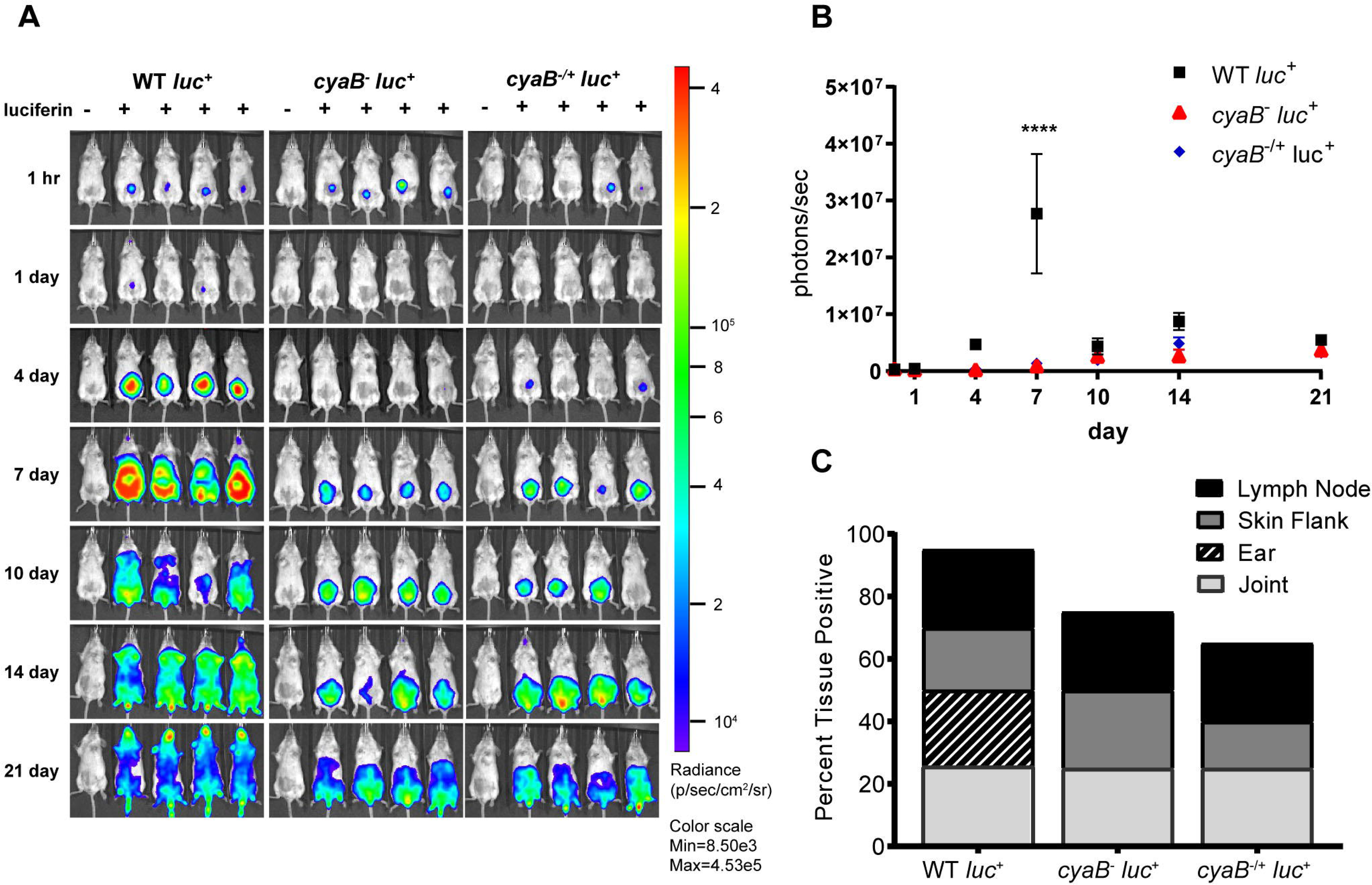
Bioluminescent *cyaB*^-^ has attenuated infection and dissemination. **(A)** Bioluminescent *B. burgdorferi* is temporal and spatial tracked during infection of Balb/c mice with 10^5^ WT, *cyaB*^-^, or c-*cyaB*-SR0623. Mice were imaged at 1 hour and 1, 4, 7, 10, 14 and 21 dpi. The mouse in the first position of the image, indicated by (-), did not receive D-luciferin to serve as a background control. n of 5 for each infection group. **(B)** The bioluminescence of four mice was quantitated and averaged. Statistical analysis was performed using Two-way ANOVA with Tukey correction relative to WT, ****p-value<0.0001. **(C)** The percentage of tissues positive for *B. burgdorferi* at 21 dpi, grown in BSKII+NRS.

## Discussion

*B. burgdorferi* gene regulation is dynamic and highly responsive to changes in environmental conditions to support the necessary adaptation for traversing between the tick vector and mammalian host (Radolf et al. 2012; D. Scott Samuels and Samuels 2016). The mechanisms used by *B. burgdorferi* to directly sense environmental conditions and relay that information to alter gene regulation are poorly understood. We hypothesize that *B. burgdorferi* uses an AC (*cyaB*, *bb0723*) and possibly cAMP for the response to environmental cues and to regulate virulence determinants important for mammalian infection. ACs cyclize ATP producing cAMP which functions as a secondary messenger in eukaryotes and prokaryotes (Stanley McKnight 1991; Botsford and Harman 1992; Kamenetsky et al. 2006; McDonough and Rodriguez 2011). In bacteria, ACs and/or cAMP are responsive to a variety of environmental changes, such as carbon starvation, CO_2_ levels, bicarbonate, osmolarity, and pH (Hoffmaster and Koehler 1997; Franchini, Ihssen, and Egli 2015; Cann et al. 2003; Rebollo-Ramirez and Larrouy-Maumus 2019). cAMP is used by numerous bacterial pathogens to alter both the host and pathogen at the level of post-transcriptional regulation for signal reception, signal transduction, AC activity, virulence gene regulation, resistance to oxidative stress, and persistence (Molina-Quiroz et al. 2018). It is well documented that *B. burgdorferi* utilizes cyclic dinucleotides during the tick and mammalian stages of pathogenesis to modulate the necessary gene regulation for response to environmental pressures, therefore it is plausible it also relies on cyclic nucleotides for regulation (Savage et al. 2015; Ye et al. 2014; Melissa J. Caimano et al. 2015; He et al. 2011; Zhang et al. 2018; Rogers et al. 2009). It is important to understand the strategies employed by *B. burgdorferi* to adapt to changing environmental conditions to evaluate borrelial pathogenesis in the context of mammalian infection.

In this study, a genetic approach was used to evaluate the borrelial *cyaB* contribution to the regulation of virulence determinants and mammalian infectivity. Borrelial *cyaB* has been annotated as a class IV AC, which is the smallest of the classes and has been previously crystalized in *Yersinia pestis* (Casjens et al. 2000; Khajanchi et al. 2016). *cyaB* is encoded on the positive strand and overlaps with *bb0722* encoded on the opposite strand. Deletion mutants of *cyaB* in two independent *B. burgdorferi* strains, 5A4-NP1 and ML23, were generated that also disrupted SR0623. Three unique complement strains, *cyaB* only, *cyaB* with SR0623, and *cyaB* with a mutagenized SR0623, were generated to clarify the contribution of *cyaB*, SR0623, and the combination of *cyaB* and SR0623 to our readouts of borrelial infectivity (Figure 1). The complete deletion of the *cyaB* ORF also truncates SR0623 and results in a reduction in the production of important mammalian virulence determinants BosR, OspC, and DbpA, while tick virulence determinants are unchanged (Figure 5). Interestingly, only *ospC* was transcriptionally down regulated in the *cyaB* deletion strains (Figure 4). Complement strains encoding the *cyaB* ORF with a truncated or mutagenized SR0623 were able to restore protein production to WT levels, suggesting the sRNA is not necessary for regulation of OspC, BosR or DbpA. Unexpectedly, the complement with *cyaB* and a complete SR0623 produced only slightly higher levels of BosR, OspC, and DbpA than the mutant. Northern analysis demonstrated higher levels of *cyaB* transcript and SR0623 in the *cyaB*-SR0623 complement strain relative to WT, which may explain the partial complementation phenotype (Figure 2). This data demonstrates *cyaB* contributes to transcriptional and post-transcriptional regulation of selected *B. burgdorferi* genes. Furthermore, *cyaB,* and possibly cAMP, are involved in regulation of factors specific for borrelial pathogenicity.

*B. burgdorferi* is greatly influenced by environmental conditions and may use *cyaB* as an environmental sensor (Radolf et al. 2012; D. Scott Samuels and Samuels 2016). We examined the *cyaB* mutant and complement strains under a variety of growth conditions by imposing oxidative stress, as well as shifting temperature, pH, and CO_2_, and found no phenotypic differences (Figure 3 & data not shown). Knowing that different carbon sources can alter cyclase activity and production of cyclic nucleotide and di-nucleotides media that replaced glucose with glycerol was used to examine regulation of borrelial virulence factors (Liu et al. 2020; Peterkofsky and Gazdar 1974). Differences in borrelial virulence determinant protein production were more pronounced in BSK-lite, independent of glycerol supplementation, relative to conventional BSKII (Figure S2 & data not shown). Further investigation indicated BSK-lite with or without glycerol resulted in the same pattern of protein production signifying glucose as the carbon source was responsible for the differential response and demonstrating borrelial catabolite repression.

*B. burgdorferi* is highly responsive to host specific signals and it is possible *cyaB* is involved in signaling for mammalian adaptation. We evaluated *cyaB* expression when co-cultured with vector or mammalian cells to mimic interactions during the pathogenic cycle. The expression of *cyaB* is induced when co-cultured with mammalian H4 neuroglial cells, but unchanged with tick ISE6 cells (Figure 6). Due to *cyaB* induced expression in the presence of mammalian cells, we also evaluated the contribution of *cyaB* to mammalian infection. During mouse infection the *cyaB* mutant strains had lower borrelial colonization, particularly during early time points (Figure 7 & 8). In later infection disseminated tissues, notably the ear and tibiotarsal joint, also had lower *B. burgdorferi* load relative to the infectious parent strain. Unfortunately, the complement strains that restored virulence determinant production in vitro did not colonize tissues at the same level as parental *B. burgdorferi.* The data is strengthened by similar results in two independent borrelial strains. Infectivity studies demonstrated that the absence of *cyaB* results in inhibited dissemination and attenuated infection. Together, this suggests *cyaB* contributes to mammalian colonization and supports this stage of the life cycle.

Khajanchi et al. evaluated the functional ability of *cyaB* and contribution to mouse infection using the *cyaB* transposon mutants (Khajanchi et al. 2016). This study showed recombinant CyaB produced cAMP in a temperature dependent manner but was not evaluated directly in *B. burgdorferi.* Our attempts to measure cAMP production during cultivation of *B. burgdorferi* have been unsuccessful likely due to the low-level expression and production of *cyaB* under these conditions. Transposon mutants with insertions in *cyaB* (insertion ratios 0.02 and 0.93) did not demonstrate an infectivity phenotype by needle inoculation or tick transmission (Khajanchi et al. 2016). Infectivity was qualitatively evaluated by borrelial outgrowth from infected tissues following a 28-day infection in which all tissues were positive for the presence of the bacteria. Our findings observed *B. burgdorferi* in most tissues at the last time point with the exception of the ears that was not assessed in the prior work. We quantitated the borrelial load of the whole mouse and individual tissues by bioluminescent imaging and qPCR, respectively. We found that overall borrelial load was reduced that were specifically lower in the skin flank and bladders during early infection, but remained low in the ear and the tibiotarsal joints. We have also investigated the involvement of SR0623 that was unknown at the time of previous studies. This indicates that *cyaB* may contribute to pathogenesis during earlier borrelial infection that can be overcome by compensatory, but unknown genes, to be able to reach a fully disseminated infection. Our study further pursued the role of borrelial ACs by investigating the regulatory effect of *cyaB* on known borrelial virulence determinants. While this study focused on a few targets it is likely that *cyaB* and cAMP have a broader impact on transcriptional and post-transcriptional regulation in *B. burgdorferi*.

Another important aspect of bacterial post-transcriptional regulation is the contribution by sRNAs. Over a thousand sRNAs were recently identified in *B. burgdorferi* (Popitsch et al. 2017), but few have been characterize. SR0623 in an intragenic sRNA that is encoded on the negative strand within the 3’ end of *cyaB* (*bb0723*) and overlaps with the hypothetical gene *bb0722* (Figure 1). SR0623 is predicted to be transcribed with *cyaB* and processed, which could result in a truncated *cyaB* transcript. Intragenic sRNAs in other bacteria regulate the genes they are encoded within, therefore SR0623 may regulate *cyaB* and/or *bb0722*. In addition, intragenic RNAs have been co-immunoprecipitated with RNA binding proteins and other mRNAs and sRNAs, indicating they may have multiple targets beside the genes they are encoded within (Melamed et al. 2020; Iosub et al. 2020; Melamed et al. 2016; Bilusic et al. 2014). In the complement with truncated SR0623 there is lower steady-state levels of *cyaB* compared to the WT, indicating SR0623 and/or the 3’ end of the *cyaB* transcript are important for regulating *cyaB* steady-state transcript levels. Furthermore, in the *cyaB* full-length SR0623 compliment there is higher steady state levels of *cyaB* and SR0623. Interestingly, both ACs and sRNAs function in post-transcriptional regulation. Finally, we also cannot rule out that our observed phenotype and challenges with complementation may be in part attributed to SR0623 regulation of *bb0722* or the intra RNA SR0622 encoded with in it.

The current study does not address the direct detection of cAMP from cultivated *B. burgdorferi.* Future studies will investigate the environmental conditions in vitro and in vivo that promotes the production of cAMP and determine if it correlates with expression or production of *cyaB*. Here in, we narrowly focused on the regulation of known tick and mammalian virulence determinant, but it is likely that *cyaB* and cAMP have a broader post-transcription regulatory impact on *B. burgdorferi.* The various strategies used to complement *cyaB* and/or SR0623 resulted in restoration phenotypes under in vitro conditions, but unfortunately were not able to completely restore the WT phenotype during mouse infection. This could be due to altered processing or stability of SR0623 and/or *cyaB* in the complement strains, which may impact the AC activity specifically induced under host adapted conditions. Our study is not the first to have difficulty complementing a borrelial gene or sRNA and represents a broader challenge in the field of bacterial pathogenesis. Further study is required to distinguish the function of intergenic sRNA from the gene it is encoded within to fully understand the complex regulation of *B. burgdorferi*.

In this study, we identified the ability of *cyaB* to contribute to the regulation of mammalian virulence determinants and infectivity in mice. This phenotype is presumably due to the production of cAMP and its impact on post-transcriptional regulation in *B. burgdorferi.* It also shed light on the complexities and possible contribution of sRNA to borrelial regulation in which the distinct responses are observed under cultivation conditions and/or during infection. It has become clear that *B. burgdorferi* utilizes post-transcriptional regulation to support pathogenesis and to provide a dynamic means to adapt to the various milieus that Lyme spirochetes move between during its complex lifecycle.

## Acknowledgements

Thank you to Dr. Jon Skare for the BosR antibody, Dr. Magnus Hook for the DbpA antibody, Dr. Richard Marconi for the Rrp2 and OspC antibodies, Dr. Seshu Janakiram for the BadR antibody, and Dr. Jenifer Coburn for P66 antibody. Appreciation to Dr. Rajesh Miranda for use of his ViiA7 Real-Time PCR machine (Applied Biosystems). We thank Dr. Ulrike Munderloh for providing ISE6 cells. We also thank Dr. Jon Skare for thoughtful comments in the editing of this manuscript.

## Author contributions

VA and JH were involved in the experimental design, data analysis and interpretation, and wrote manuscript; VA executed a majority of the experiments; LV mammalian tissue culture experiments; ES generated constructs and strains; AH & ML northern blots; ML consultation, experimental design, data analysis and interpretation; NO & RM antibody design and generation; AC tick tissue experiments. All authors were involved in manuscript editing.

---

**Supplemental Figure 1. SR0623 wobble mutation sequence.** Site directed mutagenesis of every third base pair of the sRNA SR0623 sequence is denoted by underlining. The *cyaB* ORF stop codon is indicated by an asterisk. The stop codon for the overlapping *bb0722* ORF is outlined by a box. The numbers indicate the distance from the *cyaB* ORF start codon.

**Supplemental Figure 2. Glycerol does not alter *B. burgdorferi* virulence factor production.** The *B. burgdorferi* strains 5A4-NP1 (WT) and JH522 (*cyaB*^-^) were grown in BSK-lite with or without 0.6% glycerol to mid-logarithmic phase at 32°C 1% CO_2_. Protein was harvested and resolved on a SDS-PAGE with each lane containing approximately 4×10^7^ *B. burgdorferi*. Immunoblotting was carried out using the anti-serum depicted. FlaB was used as a loading control. Representative of at least three independent replicates.

**Supplemental Figure 3. Mammalian virulence factors production in bioluminescent *cyaB* mutant.** *B. burgdorferi* ML23 strains were grown in BSK-glycerol to mid-log phase at 32°C 1% CO_2_. Protein was harvested and resolved on a SDS-PAGE with each lane containing approximately 4×10^7^ *B. burgdorferi*. Immunoblots were probed using the anti-serum depicted. FlaB was used as a loading control. Representative of at least three individual replicates. The following abbreviations are used to indicate strains: ML23 pBBE22*luc* (P), JH441 pBBE22*luc* (M), JH446 pBBE22*luc* (C).

**Supplemental Table 1.**
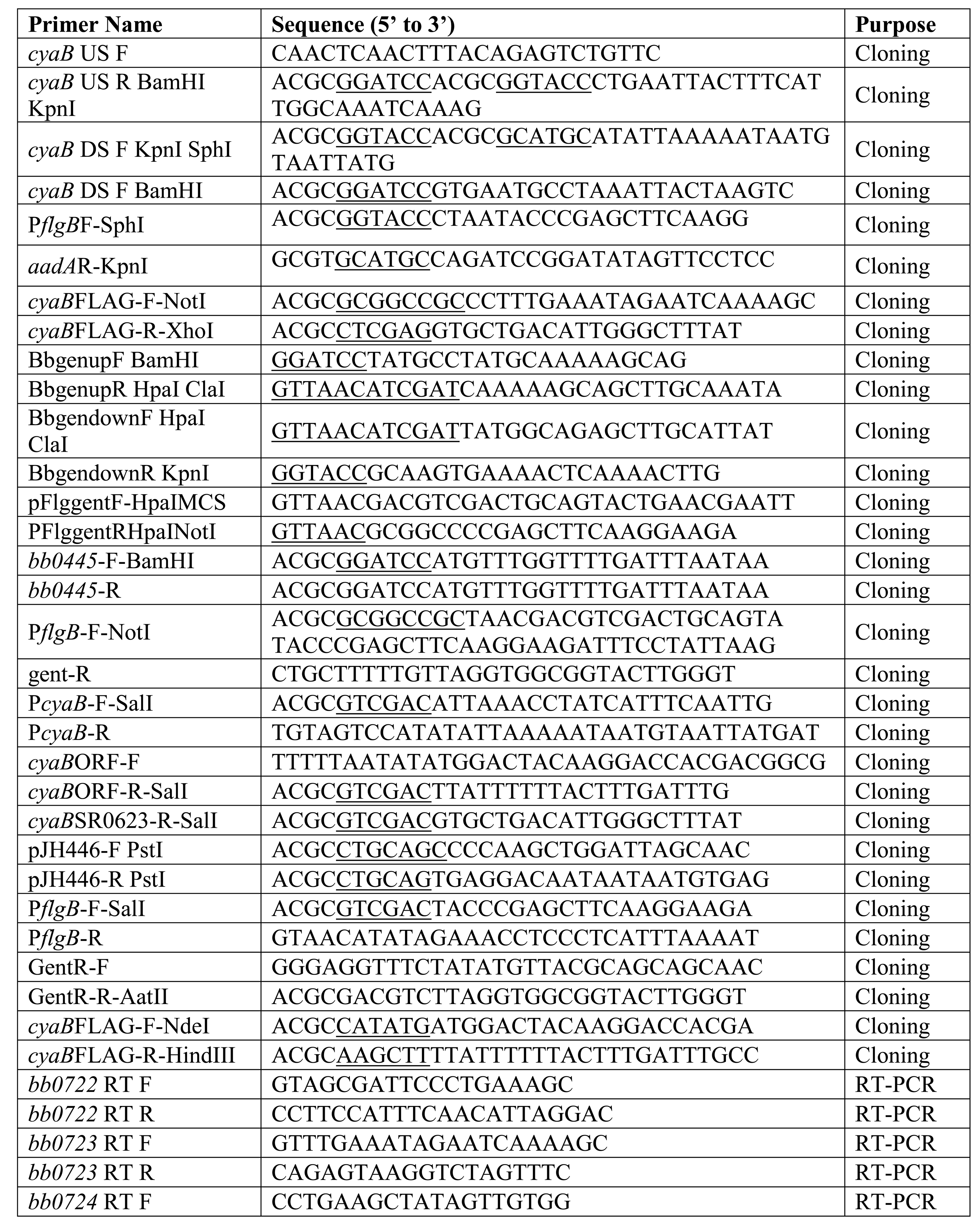
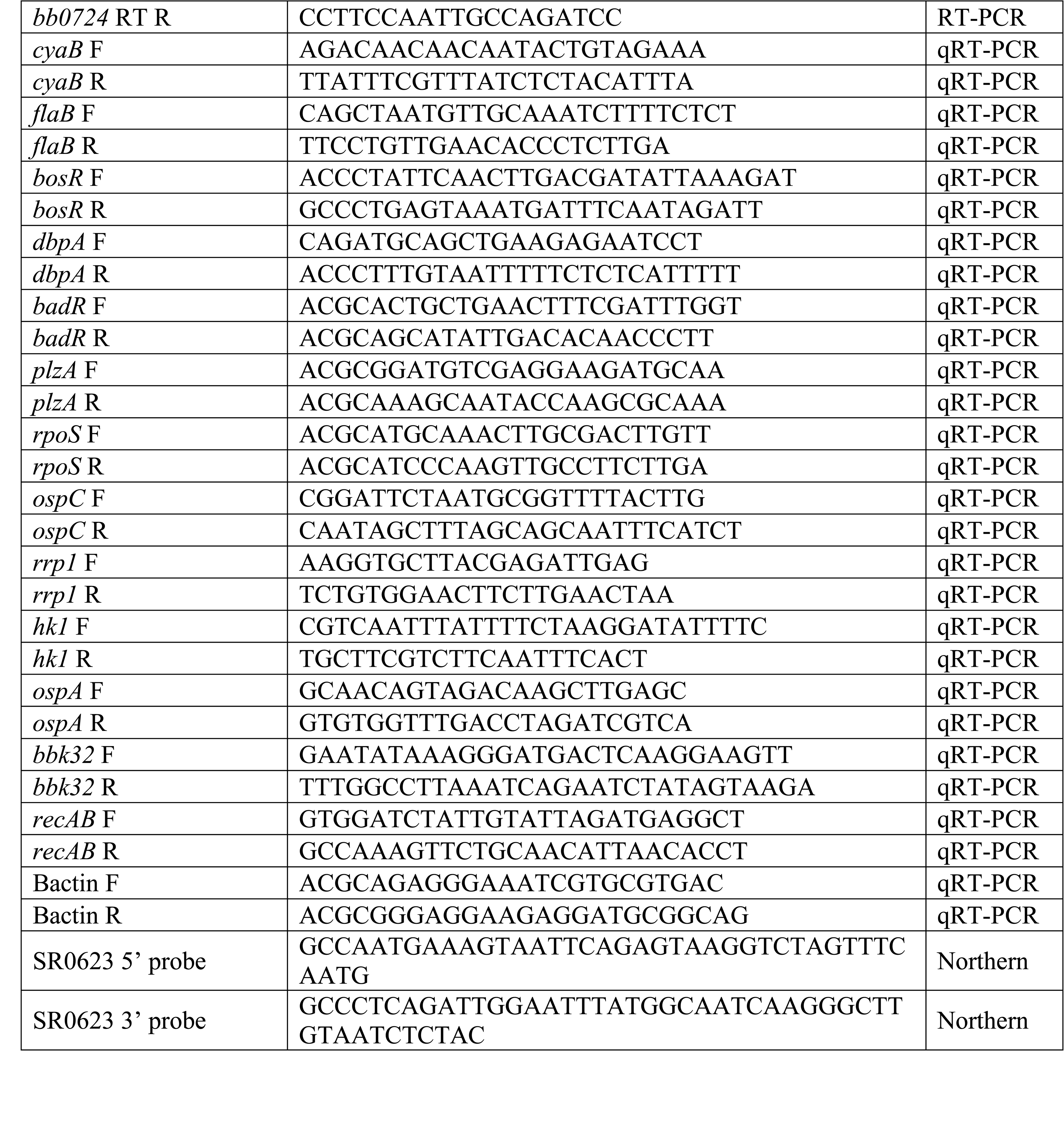
Primers used in this study.

## References

Adams, Philip P., Carlos Flores Avile, Niko Popitsch, Ivana Bilusic, Renée Schroeder, Meghan Lybecker, and Mollie W. Jewett. 2017. “In Vivo Expression Technology and 5’ End Mapping of the Borrelia Burgdorferi Transcriptome Identify Novel RNAs Expressed during Mammalian Infection.” Nucleic Acids Research 45 (2): 775–92. https://doi.org/10.1093/nar/gkw1180.

Babitzke, Paul, Ying-Jung Lai, Andrew J. Renda, and Tony Romeo. 2019. “Posttranscription Initiation Control of Gene Expression Mediated by Bacterial RNA-Binding Proteins.” Annual Review of Microbiology 73 (1): 43–67. https://doi.org/10.1146/annurev-micro-020518-115907.

Barbour, A. G. 1984. “Isolation and Cultivation of Lyme Disease Spirochetes.” The Yale Journal of Biology and Medicine 57 (4): 521–25.

Bilusic, Ivana, Niko Popitsch, Philipp Rescheneder, Renée Schroeder, and Meghan Lybecker. 2014. “Revisiting the Coding Potential of the E. Coli Genome through Hfq Co-Immunoprecipitation.” RNA Biology 11 (5): 641–54. https://doi.org/10.4161/rna.29299.

Blevins, J. S., A. T. Revel, A. H. Smith, G. N. Bachlani, and M. V. Norgard. 2007. “Adaptation of a Luciferase Gene Reporter and Lac Expression System to Borrelia Burgdorferi.” Appl Environ Microbiol 73 (5): 1501–13.

Blevins, Jon S., Haijun Xu, Ming He, Michael V. Norgard, Larry Reitzer, and X. Frank Yang. 2009. “Rrp2, a Σ54-Dependent Transcriptional Activator of Borrelia Burgdorferi, Activates RpoS in an Enhancer-Independent Manner.” Journal of Bacteriology 191 (8): 2902–5. https://doi.org/10.1128/JB.01721-08.

Bontemps-Gallo, Sébastien, Kevin Lawrence, and Frank C. Gherardini. 2016. “Two Different Virulence-Related Regulatory Pathways in Borrelia Burgdorferi Are Directly Affected by Osmotic Fluxes in the Blood Meal of Feeding Ixodes Ticks.” PLoS Pathogens 12 (8): e1005791. https://doi.org/10.1371/journal.ppat.1005791.

Botsford, J. L., and J. G. Harman. 1992. “Cyclic AMP in Prokaryotes.” Microbiology and Molecular Biology Reviews 56 (1): 100–122.

Boylan, J. A., C. S. Hummel, S. Benoit, J. Garcia-Lara, J. Treglown-Downey, E. J. Crane, and F. C. Gherardini. 2006. “Borrelia Burgdorferi Bb0728 Encodes a Coenzyme A Disulphide Reductase Whose Function Suggests a Role in Intracellular Redox and the Oxidative Stress Response.” Mol Microbiol 59 (2): 475–86.

Boylan, J. A., K. A. Lawrence, J. S. Downey, and F. C. Gherardini. 2008. “Borrelia Burgdorferi Membranes Are the Primary Targets of Reactive Oxygen Species.” Mol Microbiol 68 (3): 786–99.

Boylan, J. A., J. E. Posey, and F. C. Gherardini. 2003. “Borrelia Oxidative Stress Response Regulator, BosR: A Distinctive Zn-Dependent Transcriptional Activator.” Proc Natl Acad Sci U S A 100 (20): 11684–89.

Caimano, M. J., C. H. Eggers, K. R. Hazlett, and J. D. Radolf. 2004. “RpoS Is Not Central to the General Stress Response in Borrelia Burgdorferi but Does Control Expression of One or More Essential Virulence Determinants.” Infect Immun 72 (11): 6433–45.

Caimano, M. J., R. Iyer, C. H. Eggers, C. Gonzalez, E. A. Morton, M. A. Gilbert, I. Schwartz, and J. D. Radolf. 2007. “Analysis of the RpoS Regulon in Borrelia Burgdorferi in Response to Mammalian Host Signals Provides Insight into RpoS Function during the Enzootic Cycle.” Mol Microbiol 65 (5): 1193–1217.

Caimano, Melissa J., Star Dunham-Ems, Anna M. Allard, Maria B. Cassera, Melisha Kenedy, and Justin D. Radolf. 2015. “C-Di-GMP Modulates Gene Expression in Lyme Disease Spirochetes at the Tick-Mammal Interface to Promote Spirochete Survival during the Blood Meal and Tick-to-Mammal Transmission.” Infection and Immunity, May. https://doi.org/10.1128/IAI.00315-15.

Caimano, Melissa J., Christian H. Eggers, Cynthia A. Gonzalez, and Justin D. Radolf. 2005. “Alternate Sigma Factor RpoS Is Required for the In Vivo-Specific Repression of Borrelia Burgdorferi Plasmid Lp54-Borne OspA and Lp6.6 Genes.” Journal of Bacteriology 187 (22): 7845–52. https://doi.org/10.1128/JB.187.22.7845-7852.2005.

Caimano, Melissa J., Ashley M. Groshong, Alexia Belperron, Jialing Mao, Kelly L. Hawley, Amit Luthra, Danielle E. Graham, et al. 2019. “The RpoS Gatekeeper in Borrelia Burgdorferi: An Invariant Regulatory Scheme That Promotes Spirochete Persistence in Reservoir Hosts and Niche Diversity.” Frontiers in Microbiology 10. https://doi.org/10.3389/fmicb.2019.01923.

Caimano, Melissa J., Radha Iyer, Christian H. Eggers, Cynthia Gonzalez, Elizabeth A. Morton, Michael A. Gilbert, Ira Schwartz, and Justin D. Radolf. 2007. “Analysis of the RpoS Regulon in Borrelia Burgdorferi in Response to Mammalian Host Signals Provides Insight into RpoS Function during the Enzootic Cycle.” Molecular Microbiology 65 (5): 1193– 1217. https://doi.org/10.1111/j.1365-2958.2007.05860.x.

Cann, Martin J., Arne Hammer, Jie Zhou, and Tobias Kanacher. 2003. “A Defined Subset of Adenylyl Cyclases Is Regulated by Bicarbonate Ion*.” Journal of Biological Chemistry 278 (37): 35033–38. https://doi.org/10.1074/jbc.M303025200.

Carroll, J. A., R. M. Cordova, and C. F. Garon. 2000. “Identification of 11 PH-Regulated Genes in Borrelia Burgdorferi Localizing to Linear Plasmids.” Infect Immun 68 (12): 6677–84.

Carroll, J. A., C. F. Garon, and T. G. Schwan. 1999. “Effects of Environmental PH on Membrane Proteins in Borrelia Burgdorferi.” Infect Immun 67 (7): 3181–87.

Casjens, S., N. Palmer, R. van Vugt, W. M. Huang, B. Stevenson, P. Rosa, R. Lathigra, et al. 2000. “A Bacterial Genome in Flux: The Twelve Linear and Nine Circular Extrachromosomal DNAs in an Infectious Isolate of the Lyme Disease Spirochete Borrelia Burgdorferi.” Mol Microbiol 35 (3): 490–516.

Cugini, C., M. Medrano, T. G. Schwan, and J. Coburn. 2003. “Regulation of Expression of the Borrelia Burgdorferi Beta(3)-Chain Integrin Ligand, P66, in Ticks and in Culture.” Infect Immun 71 (2): 1001–7.

Curtiss, R., and S. M. Kelly. 1987. “Salmonella Typhimurium Deletion Mutants Lacking Adenylate Cyclase and Cyclic AMP Receptor Protein Are Avirulent and Immunogenic.” Infection and Immunity 55 (12): 3035–43.

Dong, Qing, Xufan Yan, Minhui Zheng, and Ziwen Yang. 2013. “Comparison of Two Type IV Hyperthermophilic Adenylyl Cyclases Characterizations from the Archaeon Pyrococcus Furiosus.” Journal of Molecular Catalysis B: Enzymatic 88 (April): 7–13. https://doi.org/10.1016/j.molcatb.2012.10.017.

Drecktrah, Dan, Laura S. Hall, Philipp Rescheneder, Meghan Lybecker, and D. Scott Samuels. 2018. “The Stringent Response-Regulated SRNA Transcriptome of Borrelia Burgdorferi.” Frontiers in Cellular and Infection Microbiology 8: 231. https://doi.org/10.3389/fcimb.2018.00231.

Elias, Abdallah F., James L. Bono, John J. Kupko Iii, Philip E. Stewart, Jonathan G. Krum, and Patricia A. Rosa. 2003. “New Antibiotic Resistance Cassettes Suitable for Genetic Studies in Borrelia Burgdorferi.” Journal of Molecular Microbiology and Biotechnology 6 (1): 29– 40. https://doi.org/10.1159/000073406.

Franchini, Alessandro G., Julian Ihssen, and Thomas Egli. 2015. “Effect of Global Regulators RpoS and Cyclic-AMP/CRP on the Catabolome and Transcriptome of Escherichia Coli K12 during Carbon- and Energy-Limited Growth.” PloS One 10 (7): e0133793. https://doi.org/10.1371/journal.pone.0133793.

Freedman, John C., Elizabeth A. Rogers, Jessica L. Kostick, Hongming Zhang, Radha Iyer, Ira Schwartz, and Richard T. Marconi. 2010. “Identification and Molecular Characterization of a Cyclic-Di-GMP Effector Protein, PlzA (BB0733): Additional Evidence for the Existence of a Functional Cyclic-Di-GMP Regulatory Network in the Lyme Disease Spirochete, Borrelia Burgdorferi.” FEMS Immunology and Medical Microbiology 58 (2): 285–94. https://doi.org/10.1111/j.1574-695X.2009.00635.x.

Gallagher, D. Travis, Natasha N. Smith, Sook-Kyung Kim, Annie Heroux, Howard Robinson, and Prasad T. Reddy. 2006. “Structure of the Class IV Adenylyl Cyclase Reveals a Novel Fold.” Journal of Molecular Biology 362 (1): 114–22. https://doi.org/10.1016/j.jmb.2006.07.008.

Gottesman, Susan, and Gisela Storz. 2011. “Bacterial Small RNA Regulators: Versatile Roles and Rapidly Evolving Variations.” Cold Spring Harbor Perspectives in Biology 3 (12). https://doi.org/10.1101/cshperspect.a003798.

Gstrein-Reider, E, and M Schweiger. 1982. “Regulation of Adenylate Cyclase in E. Coli.” The EMBO Journal 1 (3): 333–37.

Guo, B. P., E. L. Brown, D. W. Dorward, L. C. Rosenberg, and M. Hook. 1998. “Decorin-Binding Adhesins from Borrelia Burgdorferi.” Mol Microbiol 30 (4): 711–23.

He, Ming, Bethany K. Boardman, Dalai Yan, and X. Frank Yang. 2007. “Regulation of Expression of the Fibronectin-Binding Protein BBK32 in Borrelia Burgdorferi.” Journal of Bacteriology 189 (22): 8377–80. https://doi.org/10.1128/JB.01199-07.

He, Ming, Zhiming Ouyang, Bryan Troxell, Haijun Xu, Akira Moh, Joseph Piesman, Michael V. Norgard, Mark Gomelsky, and X. Frank Yang. 2011. “Cyclic Di-GMP Is Essential for the Survival of the Lyme Disease Spirochete in Ticks.” PLOS Pathogens 7 (6): e1002133. https://doi.org/10.1371/journal.ppat.1002133.

He, Ming, Jun-Jie Zhang, Meiping Ye, Yongliang Lou, and X. Frank Yang. 2014. “Cyclic Di-GMP Receptor PlzA Controls Virulence Gene Expression through RpoS in Borrelia Burgdorferi.” Infection and Immunity 82 (1): 445–52. https://doi.org/10.1128/IAI.01238-13.

Hoffmaster, A. R., and T. M. Koehler. 1997. “The Anthrax Toxin Activator Gene AtxA Is Associated with CO2-Enhanced Non-Toxin Gene Expression in Bacillus Anthracis.” Infect Immun 65 (8): 3091–99.

Hu, Linden T. 2016. “Lyme Disease.” Annals of Internal Medicine 165 (9): 677. https://doi.org/10.7326/L16-0409.

Hübner, A., X. Yang, D. M. Nolen, T. G. Popova, F. C. Cabello, and M. V. Norgard. 2001. “Expression of Borrelia Burgdorferi OspC and DbpA Is Controlled by a RpoN-RpoS Regulatory Pathway.” Proceedings of the National Academy of Sciences of the United States of America 98 (22): 12724–29. https://doi.org/10.1073/pnas.231442498.

Hyde, Jenny A., Dana K. Shaw, Roger Smith Iii, Jerome P. Trzeciakowski, and Jon T. Skare. 2009. “The BosR Regulatory Protein of Borrelia Burgdorferi Interfaces with the RpoS Regulatory Pathway and Modulates Both the Oxidative Stress Response and Pathogenic Properties of the Lyme Disease Spirochete.” Molecular Microbiology 74 (6): 1344–55. https://doi.org/10.1111/j.1365-2958.2009.06951.x.

Hyde, Jenny A., Dana K. Shaw, Roger Smith, Jerome P. Trzeciakowski, and Jon T. Skare. 2010. “Characterization of a Conditional BosR Mutant in Borrelia Burgdorferi” 78 (1): 265–74. https://doi.org/10.1128/IAI.01018-09.

Hyde, Jenny A., and Jon T. Skare. 2018. “Detection of Bioluminescent Borrelia Burgdorferi from In Vitro Cultivation and During Murine Infection.” Methods in Molecular Biology (Clifton, N.J.) 1690: 241–57. https://doi.org/10.1007/978-1-4939-7383-5_18.

Hyde, Jenny A., Jerome P. Trzeciakowski, and Jonathan T. Skare. 2007. “Borrelia Burgdorferi Alters Its Gene Expression and Antigenic Profile in Response to CO2 Levels.” Journal of Bacteriology 189 (2): 437–45. https://doi.org/10.1128/JB.01109-06.

Hyde, Jenny A., Eric H. Weening, MiHee Chang, Jerome P. Trzeciakowski, Magnus Höök, Jeffrey D. Cirillo, and Jon T. Skare. 2011. “Bioluminescent Imaging of Borrelia Burgdorferi in Vivo Demonstrates That the Fibronectin-Binding Protein BBK32 Is Required for Optimal Infectivity.” Molecular Microbiology 82 (1): 99–113. https://doi.org/10.1111/j.1365-2958.2011.07801.x.

Hyde, Jenny A., Eric H. Weening, and Jon T. Skare. 2011. “Genetic Transformation of Borrelia Burgdorferi.” Current Protocols in Microbiology 20 (1): 12C.4.1–12C.4.17. https://doi.org/10.1002/9780471729259.mc12c04s20.

Iosub, Ira Alexandra, Robert Willem van Nues, Stuart William McKellar, Karen Jule Nieken, Marta Marchioretto, Brandon Sy, Jai Justin Tree, Gabriella Viero, and Sander Granneman. 2020. “Hfq CLASH Uncovers SRNA-Target Interaction Networks Linked to Nutrient Availability Adaptation.” Edited by Joseph T Wade, James L Manley, and Ben F Luisi. ELife 9 (May): e54655. https://doi.org/10.7554/eLife.54655.

Izac, Jerilyn R., Andrew C. Camire, Christopher G. Earnhart, Monica E. Embers, Rebecca A. Funk, Edward B. Breitschwerdt, and Richard T. Marconi. 2019. “Analysis of the Antigenic Determinants of the OspC Protein of the Lyme Disease Spirochetes: Evidence That the C10 Motif Is Not Immunodominant or Required to Elicit Bactericidal Antibody Responses.” Vaccine 37 (17): 2401–7. https://doi.org/10.1016/j.vaccine.2019.02.007.

Kamenetsky, Margarita, Sabine Middelhaufe, Erin M. Bank, Lonny R. Levin, Jochen Buck, and Clemens Steegborn. 2006. “Molecular Details of CAMP Generation in Mammalian Cells: A Tale of Two Systems.” Journal of Molecular Biology 362 (4): 623–39. https://doi.org/10.1016/j.jmb.2006.07.045.

Kawabata, H., S. J. Norris, and H. Watanabe. 2004. “BBE02 Disruption Mutants of Borrelia Burgdorferi B31 Have a Highly Transformable, Infectious Phenotype.” Infect Immun 72 (12): 7147–54.

Khajanchi, Bijay K., Evelyn Odeh, Lihui Gao, Mary B. Jacobs, Mario T. Philipp, Tao Lin, and Steven J. Norris. 2016. “Phosphoenolpyruvate Phosphotransferase System Components Modulate Gene Transcription and Virulence of Borrelia Burgdorferi.” Infection and Immunity 84 (3): 754–64. https://doi.org/10.1128/IAI.00917-15.

Konkel, M. E., and K. Tilly. 2000. “Temperature-Regulated Expression of Bacterial Virulence Genes.” Microbes Infect 2 (2): 157–66.

Kostick, Jessica L., Lee T. Szkotnicki, Elizabeth A. Rogers, Paola Bocci, Nadia Raffaelli, and Richard T. Marconi. 2011. “The Diguanylate Cyclase, Rrp1, Regulates Critical Steps in the Enzootic Cycle of the Lyme Disease Spirochetes.” Molecular Microbiology 81 (1): 219–31. https://doi.org/10.1111/j.1365-2958.2011.07687.x.

Kostick-Dunn, Jessica L., Jerilyn R. Izac, John C. Freedman, Lee T. Szkotnicki, Lee D. Oliver, and Richard T. Marconi. 2018. “The Borrelia Burgdorferi C-Di-GMP Binding Receptors, PlzA and PlzB, Are Functionally Distinct.” Frontiers in Cellular and Infection Microbiology 8: 213. https://doi.org/10.3389/fcimb.2018.00213.

Labandeira-Rey, M., and J. T. Skare. 2001. “Decreased Infectivity in Borrelia Burgdorferi Strain B31 Is Associated with Loss of Linear Plasmid 25 or 28-1.” Infection and Immunity 69 (1): 446–55. https://doi.org/10.1128/IAI.69.1.446-455.2001.

Lackum, Kate von, and Brian Stevenson. 2005. “Carbohydrate Utilization by the Lyme Borreliosis Spirochete, Borrelia Burgdorferi.” FEMS Microbiology Letters 243 (1): 173–79. https://doi.org/10.1016/j.femsle.2004.12.002.

Lawrenz, M. B., H. Kawabata, J. E. Purser, and S. J. Norris. 2002. “Decreased Electroporation Efficiency in Borrelia Burgdorferi Containing Linear Plasmids Lp25 and Lp56: Impact on Transformation of Infectious B. Burgdorferi.” Infect Immun 70 (9): 4798–4804.

Li, X., U. Pal, N. Ramamoorthi, X. Liu, D. C. Desrosiers, C. H. Eggers, J. F. Anderson, J. D. Radolf, and E. Fikrig. 2007. “The Lyme Disease Agent Borrelia Burgdorferi Requires BB0690, a Dps Homologue, to Persist within Ticks.” Mol Microbiol 63 (3): 694–710.

Liang, Weili, Alberto Pascual-Montano, Anisia J. Silva, and Jorge A. Benitez. 2007. “The Cyclic AMP Receptor Protein Modulates Quorum Sensing, Motility and Multiple Genes That Affect Intestinal Colonization in Vibrio Cholerae.” Microbiology, 153 (9): 2964–75. https://doi.org/10.1099/mic.0.2007/006668-0.

Liu, Cong, Di Sun, Jingrong Zhu, Jiawen Liu, and Weijie Liu. 2020. “The Regulation of Bacterial Biofilm Formation by CAMP-CRP: A Mini-Review.” Frontiers in Microbiology 11. https://doi.org/10.3389/fmicb.2020.00802.

Livak, K. J., and T. D. Schmittgen. 2001. “Analysis of Relative Gene Expression Data Using Real-Time Quantitative PCR and the 2(-Delta Delta C(T)) Method.” Methods (San Diego, Calif.) 25 (4): 402–8. https://doi.org/10.1006/meth.2001.1262.

Livengood, Jill A., Virginia L. Schmit, and Robert D. Gilmore. 2008. “Global Transcriptome Analysis of Borrelia Burgdorferi during Association with Human Neuroglial Cells.” Infection and Immunity 76 (1): 298–307. https://doi.org/10.1128/IAI.00866-07.

Lybecker, M., and KC Henderson. 2018. “Borrelia Burgdorferi Transcriptome Analysis by RNA-Sequencing.” In Borrelia Burgdorferi: Methods and Protocols, edited by Utpal Pal and Ozlem Buyuktanir, 127–36. Methods in Molecular Biology. New York, NY: Springer.

Lybecker, Meghan, Bob Zimmermann, Ivana Bilusic, Nadezda Tukhtubaeva, and Renée Schroeder. 2014. “The Double-Stranded Transcriptome of Escherichia Coli.” Proceedings of the National Academy of Sciences of the United States of America 111 (8): 3134–39. https://doi.org/10.1073/pnas.1315974111.

Mallory, Katherine L., Daniel P. Miller, Lee D. Oliver, John C. Freedman, Jessica L. Kostick-Dunn, Jason A. Carlyon, James D. Marion, Jessica K. Bell, and Richard T. Marconi. 2016. “Cyclic-Di-GMP Binding Induces Structural Rearrangements in the PlzA and PlzC Proteins of the Lyme Disease and Relapsing Fever Spirochetes: A Possible Switch Mechanism for c-Di-GMP-Mediated Effector Functions.” Pathogens and Disease 74 (8). https://doi.org/10.1093/femspd/ftw105.

Maruskova, Mahulena, M. Dolores Esteve-Gassent, Valerie L. Sexton, and J. Seshu. 2008. “Role of the BBA64 Locus of Borrelia Burgdorferi in Early Stages of Infectivity in a Murine Model of Lyme Disease.” Infection and Immunity 76 (1): 391–402. https://doi.org/10.1128/IAI.01118-07.

McDonough, Kathleen A., and Ana Rodriguez. 2011. “The Myriad Roles of Cyclic AMP in Microbial Pathogens: From Signal to Sword.” Nature Reviews Microbiology 10 (November): 27.

Melamed, Sahar, Philip P. Adams, Aixia Zhang, Hongen Zhang, and Gisela Storz. 2020. “RNA-RNA Interactomes of ProQ and Hfq Reveal Overlapping and Competing Roles.” Molecular Cell 77 (2): 411–425.e7. https://doi.org/10.1016/j.molcel.2019.10.022.

Melamed, Sahar, Asaf Peer, Raya Faigenbaum-Romm, Yair E. Gatt, Niv Reiss, Amir Bar, Yael Altuvia, Liron Argaman, and Hanah Margalit. 2016. “Global Mapping of Small RNA-Target Interactions in Bacteria.” Molecular Cell 63 (5): 884–97. https://doi.org/10.1016/j.molcel.2016.07.026.

Miller, Christine L., S. L. Rajasekhar Karna, and J. Seshu. 2013. “Borrelia Host Adaptation Regulator (BadR) Regulates RpoS to Modulate Host Adaptation and Virulence Factors in Borrelia Burgdorferi.” Molecular Microbiology 88 (1): 105–24. https://doi.org/10.1111/mmi.12171.

Miller, Daniel P., Lee D. Oliver, Brittney K. Tegels, Lucas A. Reed, Nathaniel S. O’Bier, Kurni Kurniyati, Lindsay A. Faust, et al. 2016. “The Treponema Denticola FhbB Protein Is a Dominant Early Antigen That Elicits FhbB Variant-Specific Antibodies That Block Factor H Binding and Cleavage by Dentilisin.” Infection and Immunity 84 (7): 2051–58. https://doi.org/10.1128/IAI.01542-15.

Molina-Quiroz, Roberto C., Cecilia Silva-Valenzuela, Jennifer Brewster, Eduardo Castro-Nallar, Stuart B. Levy, and Andrew Camilli. 2018. “Cyclic AMP Regulates Bacterial Persistence through Repression of the Oxidative Stress Response and SOS-Dependent DNA Repair in Uropathogenic Escherichia Coli.” MBio 9 (1). https://doi.org/10.1128/mBio.02144-17.

Mouali, Youssef El, Tania Gaviria-Cantin, María Antonia Sánchez-Romero, Marta Gibert, Alexander J. Westermann, Jörg Vogel, and Carlos Balsalobre. 2018. “CRP-CAMP Mediates Silencing of Salmonella Virulence at the Post-Transcriptional Level.” PLOS Genetics 14 (6): e1007401. https://doi.org/10.1371/journal.pgen.1007401.

Novak, Elizabeth A., Syed Z. Sultan, and Md A. Motaleb. 2014. “The Cyclic-Di-GMP Signaling Pathway in the Lyme Disease Spirochete, Borrelia Burgdorferi.” Frontiers in Cellular and Infection Microbiology 4: 56. https://doi.org/10.3389/fcimb.2014.00056.

Oliver, J. H., F. W. Chandler, M. P. Luttrell, A. M. James, D. E. Stallknecht, B. S. McGuire, H. J. Hutcheson, G. A. Cummins, and R. S. Lane. 1993. “Isolation and Transmission of the Lyme Disease Spirochete from the Southeastern United States.” Proceedings of the National Academy of Sciences of the United States of America 90 (15): 7371–75.

Oliver, Jonathan D., Adela S. Oliva Chávez, Roderick F. Felsheim, Timothy J. Kurtti, and Ulrike G. Munderloh. 2015. “An Ixodes Scapularis Cell Line with a Predominantly Neuron-like Phenotype.” Experimental & Applied Acarology 66 (3): 427–42. https://doi.org/10.1007/s10493-015-9908-1.

Ouyang, Z., J. S. Blevins, and M. V. Norgard. 2008. “Transcriptional Interplay among the Regulators Rrp2, RpoN and RpoS in Borrelia Burgdorferi.” Microbiology (Reading, England) 154 (Pt 9): 2641–58.

Ouyang, Zhiming, Ranjit K. Deka, and Michael V. Norgard. 2011. “BosR (BB0647) Controls the RpoN-RpoS Regulatory Pathway and Virulence Expression in Borrelia Burgdorferi by a Novel DNA-Binding Mechanism.” PLOS Pathogens 7 (2): e1001272. https://doi.org/10.1371/journal.ppat.1001272.

Ouyang, Zhiming, Manish Kumar, Toru Kariu, Shayma Haq, Martin Goldberg, Utpal Pal, and Michael V. Norgard. 2009. “BosR (BB0647) Governs Virulence Expression in Borrelia Burgdorferi.” Molecular Microbiology 74 (6): 1331–43. https://doi.org/10.1111/j.1365-2958.2009.06945.x.

Papenfort, Kai, and Jörg Vogel. 2010. “Regulatory RNA in Bacterial Pathogens.” Cell Host & Microbe 8 (1): 116–27. https://doi.org/10.1016/j.chom.2010.06.008.

Peterkofsky, Alan, and Celia Gazdar. 1974. “Glucose Inhibition of Adenylate Cyclase in Intact Cells of Escherichia Coli B.” Proceedings of the National Academy of Sciences of the United States of America 71 (6): 2324–28.

Popitsch, Niko, Ivana Bilusic, Philipp Rescheneder, Renée Schroeder, and Meghan Lybecker. 2017. “Temperature-Dependent SRNA Transcriptome of the Lyme Disease Spirochete.” BMC Genomics 18 (January). https://doi.org/10.1186/s12864-016-3398-3.

Purificação, Aline Dias da, Nathalia Marins de Azevedo, Gabriel Guarany de Araujo, Robson Francisco de Souza, and Cristiane Rodrigues Guzzo. 2020. “The World of Cyclic Dinucleotides in Bacterial Behavior.” Molecules 25 (10): 2462. https://doi.org/10.3390/molecules25102462.

Radolf, Justin D, Melissa J Caimano, Brian Stevenson, and Linden T Hu. 2012. “Of Ticks, Mice and Men: Understanding the Dual-Host Lifestyle of Lyme Disease Spirochaetes.” Nature Reviews. Microbiology 10 (2): 87–99. https://doi.org/10.1038/nrmicro2714.

Ramsey, Meghan E., Jenny A. Hyde, Diana N. Medina-Perez, Tao Lin, Lihui Gao, Maureen E. Lundt, Xin Li, Steven J. Norris, Jon T. Skare, and Linden T. Hu. 2017. “A High-Throughput Genetic Screen Identifies Previously Uncharacterized Borrelia Burgdorferi Genes Important for Resistance against Reactive Oxygen and Nitrogen Species.” PLOS Pathogens 13 (2): e1006225. https://doi.org/10.1371/journal.ppat.1006225.

Rebollo-Ramirez, Sonia, and Gerald Larrouy-Maumus. 2019. “NaCl Triggers the CRP-Dependent Increase of CAMP in Mycobacterium Tuberculosis.” Tuberculosis (Edinburgh, Scotland) 116: 8–16. https://doi.org/10.1016/j.tube.2019.03.009.

Rogers, Elizabeth A., Darya Terekhova, Hong-Ming Zhang, Kelley M. Hovis, Ira Schwartz, and Richard T. Marconi. 2009. “Rrp1, a Cyclic-Di-GMP-Producing Response Regulator, Is an Important Regulator of Borrelia Burgdorferi Core Cellular Functions.” Molecular Microbiology 71 (6): 1551–73. https://doi.org/10.1111/j.1365-2958.2009.06621.x.

Rosenberg, Ronald. 2018. “Vital Signs: Trends in Reported Vectorborne Disease Cases — United States and Territories, 2004–2016.” MMWR. Morbidity and Mortality Weekly Report 67. https://doi.org/10.15585/mmwr.mm6717e1.

Samuels, D S, K E Mach, and C F Garon. 1994. “Genetic Transformation of the Lyme Disease Agent Borrelia Burgdorferi with Coumarin-Resistant GyrB.” Journal of Bacteriology 176 (19): 6045–49.

Samuels, D. Scott, and Leah R. N. Samuels. 2016. “Gene Regulation During the Enzootic Cycle of the Lyme Disease Spirochete.” Forum on Immunopathological Diseases and Therapeutics 7 (3–4): 205–12. https://doi.org/10.1615/ForumImmunDisTher.2017019469.

Saputra, Elizabeth P., Jerome P. Trzeciakowski, and Jenny A. Hyde. 2020. “Borrelia Burgdorferi Spatiotemporal Regulation of Transcriptional Regulator BosR and Decorin Binding Protein during Murine Infection.” Scientific Reports 10 (1): 12534. https://doi.org/10.1038/s41598-020-69212-7.

Savage, Christina R., William K. Arnold, Alexandra Gjevre-Nail, Benjamin J. Koestler, Eric L. Bruger, Jeffrey R. Barker, Christopher M. Waters, and Brian Stevenson. 2015. “Intracellular Concentrations of Borrelia Burgdorferi Cyclic Di-AMP Are Not Changed by Altered Expression of the CdaA Synthase.” PloS One 10 (4): e0125440. https://doi.org/10.1371/journal.pone.0125440.

Schmit, Virginia L., Toni G. Patton, and Robert D. Gilmore. 2011. “Analysis of Borrelia Burgdorferi Surface Proteins as Determinants in Establishing Host Cell Interactions.” Frontiers in Microbiology 2: 141. https://doi.org/10.3389/fmicb.2011.00141.

Seshu, J., J. A. Boylan, F. C. Gherardini, and J. T. Skare. 2004. “Dissolved Oxygen Levels Alter Gene Expression and Antigen Profiles in Borrelia Burgdorferi.” Infect Immun 72 (3): 1580–86.

Seshu, J., J. A. Boylan, J. A. Hyde, K. L. Swingle, F. C. Gherardini, and J. T. Skare. 2004. “A Conservative Amino Acid Change Alters the Function of BosR, the Redox Regulator of Borrelia Burgdorferi.” Mol Microbiol 54 (5): 1352–63.

Sismeiro, O., P. Trotot, F. Biville, C. Vivares, and A. Danchin. 1998. “Aeromonas Hydrophila Adenylyl Cyclase 2: A New Class of Adenylyl Cyclases with Thermophilic Properties and Sequence Similarities to Proteins from Hyperthermophilic Archaebacteria.” Journal of Bacteriology 180 (13): 3339–44. https://doi.org/10.1128/JB.180.13.3339-3344.1998.

Smith, A. H., J. S. Blevins, G. N. Bachlani, X. F. Yang, and M. V. Norgard. 2007. “Evidence That RpoS (SigmaS) in Borrelia Burgdorferi Is Controlled Directly by RpoN (Sigma54/SigmaN).” J Bacteriol 189 (5): 2139–44.

Smith, Natasha, Sook-Kyung Kim, Prasad T. Reddy, and D. Travis Gallagher. 2006. “Crystallization of the Class IV Adenylyl Cyclase from Yersinia Pestis.” Acta Crystallographica Section F: Structural Biology and Crystallization Communications 62 (Pt 3): 200–204. https://doi.org/10.1107/S1744309106002855.

Smith, Roger S., Matthew C. Wolfgang, and Stephen Lory. 2004. “An Adenylate Cyclase-Controlled Signaling Network Regulates Pseudomonas Aeruginosa Virulence in a Mouse Model of Acute Pneumonia.” Infection and Immunity 72 (3): 1677–84. https://doi.org/10.1128/iai.72.3.1677-1684.2004.

Stanek, Gerold, and Franc Strle. 2018. “Lyme Borreliosis–from Tick Bite to Diagnosis and Treatment.” FEMS Microbiology Reviews 42 (3): 233–58. https://doi.org/10.1093/femsre/fux047.

Stanley McKnight, G. 1991. “Cyclic AMP Second Messenger Systems.” Current Opinion in Cell Biology 3 (2): 213–17. https://doi.org/10.1016/0955-0674(91)90141-K.

Steere, Allen C., Franc Strle, Gary P. Wormser, Linden T. Hu, John A. Branda, Joppe W. R. Hovius, Xin Li, and Paul S. Mead. 2016. “Lyme Borreliosis.” Nature Reviews. Disease Primers 2: 16090. https://doi.org/10.1038/nrdp.2016.90.

Stevenson, B., T. G. Schwan, and P. A. Rosa. 1995. “Temperature-Related Differential Expression of Antigens in the Lyme Disease Spirochete, Borrelia Burgdorferi.” Infection & Immunity 63 (11): 4535–39.

Sultan, Syed Z., Joshua E. Pitzer, Tristan Boquoi, Gerry Hobbs, Michael R. Miller, and M. A. Motaleb. 2011. “Analysis of the HD-GYP Domain Cyclic Dimeric GMP Phosphodiesterase Reveals a Role in Motility and the Enzootic Life Cycle of Borrelia Burgdorferi.” Infection and Immunity 79 (8): 3273–83. https://doi.org/10.1128/IAI.05153-11.

Sultan, Syed Z., Joshua E. Pitzer, Michael R. Miller, and Md A. Motaleb. 2010. “Analysis of a Borrelia Burgdorferi Phosphodiesterase Demonstrates a Role for Cyclic-Di-Guanosine Monophosphate in Motility and Virulence.” Molecular Microbiology 77 (1): 128–42. https://doi.org/10.1111/j.1365-2958.2010.07191.x.

Tokarz, Rafal, Julie M. Anderton, Laura I. Katona, and Jorge L. Benach. 2004. “Combined Effects of Blood and Temperature Shift on Borrelia Burgdorferi Gene Expression as Determined by Whole Genome DNA Array.” Infection and Immunity 72 (9): 5419–32. https://doi.org/10.1128/IAI.72.9.5419-5432.2004.

Wu, Jing, Eric H. Weening, Jennifer B. Faske, Magnus Höök, and Jon T. Skare. 2011. “Invasion of Eukaryotic Cells by Borrelia Burgdorferi Requires Β1 Integrins and Src Kinase Activity.” Infection and Immunity 79 (3): 1338–48. https://doi.org/10.1128/IAI.01188-10.

Yang, X. F., M. C. Lybecker, U. Pal, S. M. Alani, J. Blevins, A. T. Revel, D. S. Samuels, and M. V. Norgard. 2005. “Analysis of the OspC Regulatory Element Controlled by the RpoN-RpoS Regulatory Pathway in Borrelia Burgdorferi.” J Bacteriol 187 (14): 4822–29.

Yang, X., M. S. Goldberg, T. G. Popova, G. B. Schoeler, S. K. Wikel, K. E. Hagman, and M. V. Norgard. 2000. “Interdependence of Environmental Factors Influencing Reciprocal Patterns of Gene Expression in Virulent Borrelia Burgdorferi.” Mol Microbiol 37 (6): 1470–9.

Yang, Xiaofeng F., Sophie M. Alani, and Michael V. Norgard. 2003. “The Response Regulator Rrp2 Is Essential for the Expression of Major Membrane Lipoproteins in Borrelia Burgdorferi.” Proceedings of the National Academy of Sciences 100 (19): 11001–6. https://doi.org/10.1073/pnas.1834315100.

Ye, Meiping, Jun-Jie Zhang, Xin Fang, Gavin B. Lawlis, Bryan Troxell, Yan Zhou, Mark Gomelsky, Yongliang Lou, and X. Frank Yang. 2014. “DhhP, a Cyclic Di-AMP Phosphodiesterase of Borrelia Burgdorferi, Is Essential for Cell Growth and Virulence.” Infection and Immunity 82 (5): 1840–49. https://doi.org/10.1128/IAI.00030-14.

Yin, Wen, Xia Cai, Hongdan Ma, Li Zhu, Yuling Zhang, Shan-Ho Chou, Michael Y Galperin, and Jin He. 2020. “A Decade of Research on the Second Messenger C-Di-AMP.” FEMS Microbiology Reviews 44 (6): 701–24. https://doi.org/10.1093/femsre/fuaa019.

Zhang, Jun-Jie, Tong Chen, Youyun Yang, Jimei Du, Hongxia Li, Bryan Troxell, Ming He, Sebastian E. Carrasco, Mark Gomelsky, and X. Frank Yang. 2018. “Positive and Negative Regulation of Glycerol Utilization by the C-Di-GMP Binding Protein PlzA in Borrelia Burgdorferi.” Journal of Bacteriology 200 (22). https://doi.org/10.1128/JB.00243-18.

