## Supplementary figures and images for "The *Borrelia burgdorferi* adenylyl cyclase, CyaB, is important for virulence factor production and mammalian infection"

### Supplemental Figure 1

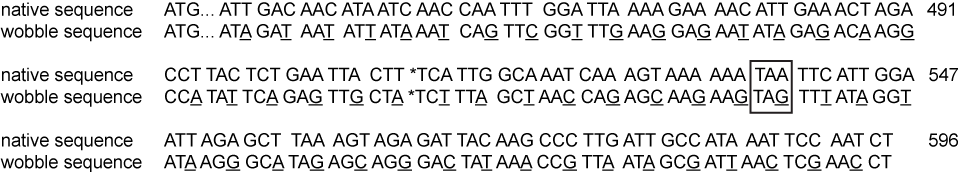

### Supplemental Figure 2

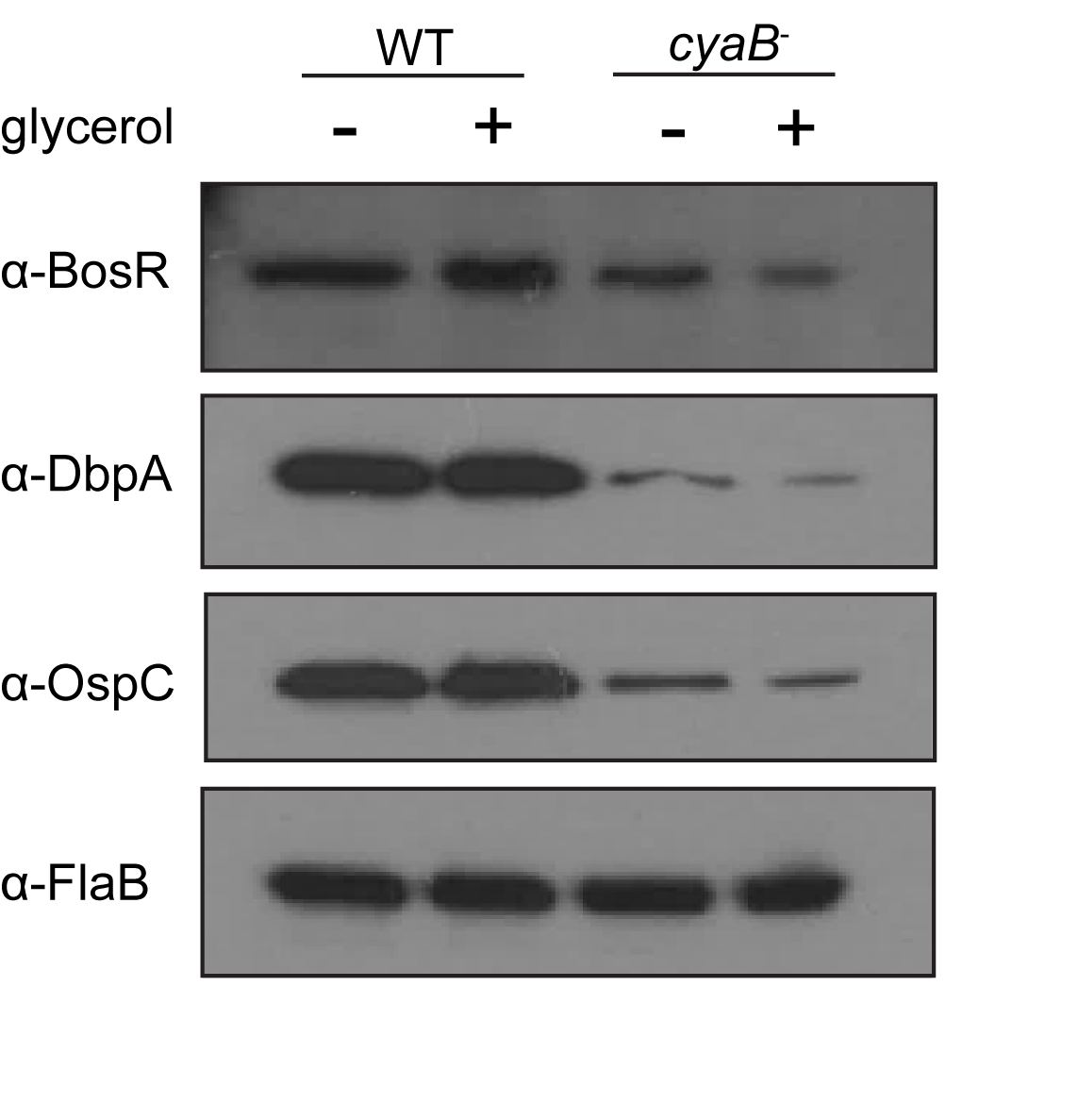

### Supplemental Figure 3

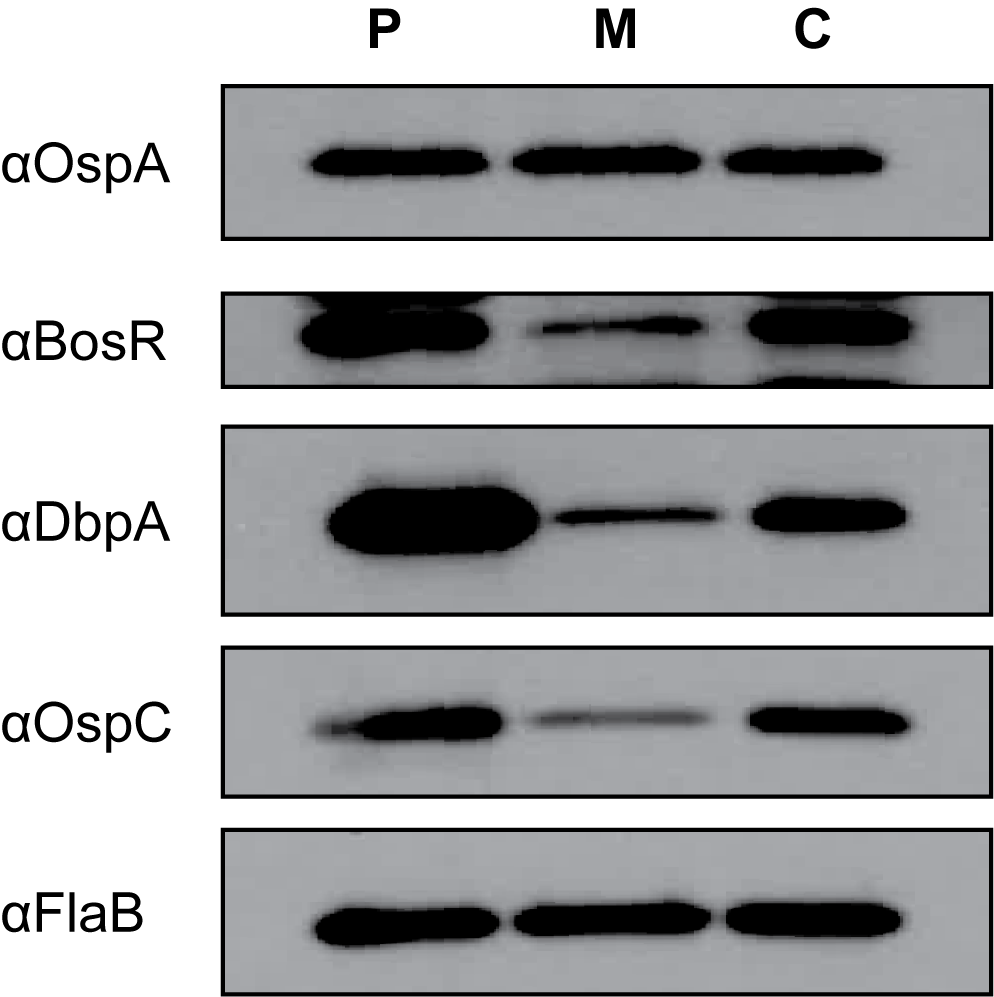
